# Unravelling genome-wide mosaic microsatellite mutations at single-cell resolution

**DOI:** 10.64898/2026.02.04.703915

**Authors:** Chunyi Wang, Wenxuan Fan, Weixiang Wang, Yonghe Xia, Jinhong Lu, Xiaoyu Ma, Jichuan Yu, Yunchao Zheng, Qing Yang, Meizhen Lin, Xiuyan Yu, Xushen Xiong, Manolis Kellis, Qi Xie, Yanmei Dou, Data Group

## Abstract

Short tandem repeats (STRs), or microsatellites, are highly mutable genomic elements that modulate gene regulations and are implicated in a range of human diseases. However, detecting mosaic STR mutations at single-cell resolution remains challenging due to both technical and biological complexities. To address this, we developed BayesMonSTR, a robust algorithm that enables accurate detection of mosaic STR mutations. Using this tool in single-cell analysis of human tissues, we reveal an accumulation of longer mosaic STR insertions and deletions (indels) in aging mitotic and post-mitotic cells. Strikingly, prefrontal cortex (PFC) neurons accumulate a higher burden of STR mutations than B cells or lung epithelium, with aged neurons exhibiting a particularly pronounced increase in longer STR deletions. These mutations are enriched at transcription start sites (TSSs) and active enhancers of highly expressed genes. Our work establishes a foundation for genome-wide, hypothesis-free discovery of disease-associated mosaic STR mutations and reveals a previously unexplored landscape of mosaic STR variation in development and aging.

---

Somatic mosaicism, arising from postzygotic mutations accumulated throughout life, represents the dynamic interface between genome instability, environmental exposures, and cellular repair processes, and is a recognized contributor to aging and disease^1^. STR regions are vastly overrepresented in the human genome^2^, and their intrinsic hypermutability—occurring at a rate roughly 100 times that of non-repeat regions^3^— renders them particularly vulnerable to mutations. Although STRs play essential roles in regulating transcription, facilitating chromatin interactions, and are linked to nearly a hundred human disorders^4,5^, mosaic STR mutations in aging tissues remain largely unexplored. Current genotyping methods^6–8^ are unable to resolve mosaic STR mutations at single-cell resolution, constrained by both technical challenges (e.g., amplification & sequence errors, alignment ambiguities) and the inherent biological complexity of repetitive regions (e.g., repeat sequence interruptions, truncated motifs).

Recognizing this critical gap, we developed BayesMonSTR—a computational framework that integrates probabilistic read segmentation, haplotype phasing, a hierarchical Bayesian network, and a machine learning classification model to detect mosaic STR insertions, deletions (indels) and mismatches from single cells with nucleotide-level resolution (Fig. 1a-f, Supplementary Fig. 1, Supplementary Table 1, Methods). By integrating multiple layers of data—at the population, individual, and single-cell levels—and probabilistically quantifying all aspects (e.g., sequence error rate, read phasing probability, repeat segmentation, and genotyping likelihoods), BayesMonSTR resolves ambiguities in both extracting imperfect STR alleles from noisy data and robust detection of mutation candidates. Furthermore, it rigorously filters out technical artifacts and germline polymorphisms using a machine learning model trained based on a multifaceted set of “gold-standard” orthogonal benchmarking data generated in this study (Fig. 1g-h; see Supplementary Information for details on the validation strategies: duplicate single-cell libraries facilitated by single-cell DNA & RNA cross validation, and deep bulk DNA sequencing from adjacent tissues).

**Fig. 1.**
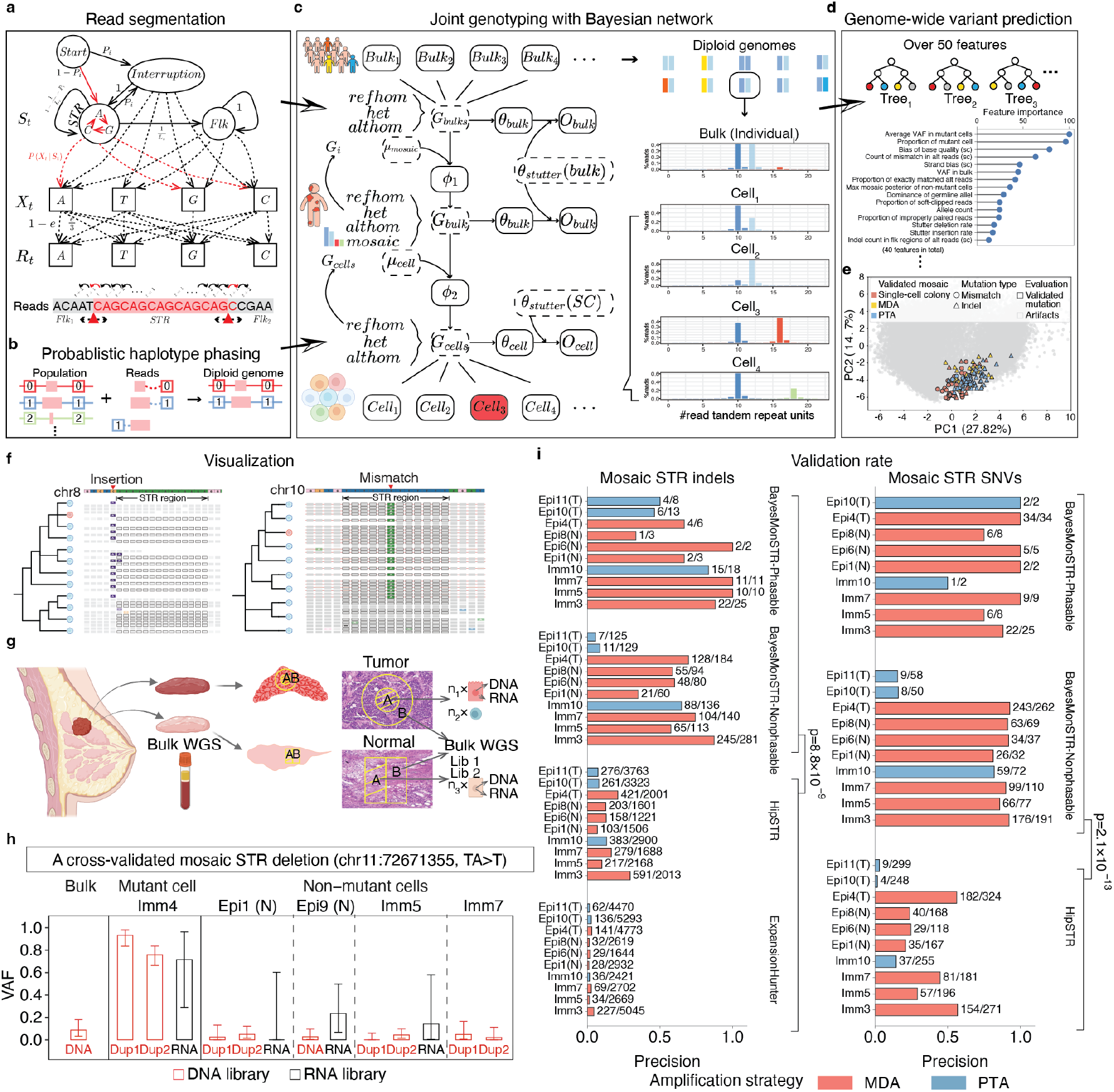
Algorithm and benchmark of BayesMonSTR. (**a**) BayesMonSTR employs an HMM to probabilistically segment each sequencing read, distinguishing the STR region from its flanking sequences. (**b**) STR alleles are assigned to their chromosomal haplotypes by statistically linking them to the alleles of neighboring heterozygous variants. (**c**) A Hierarchical Bayesian Network integrates three layers of data— population-level sequencing, individual-level bulk sequencing, and single cell sequencing—to calculate individual- and cell-level genotype posteriors. (**d**) A random forest model, trained on over 50 carefully chosen read-level and genome-level features, classifies true mosaic mutations from artifacts. (**e**) PCA constructed from top 40 features shows that validated mosaic variants from independent single-cell datasets cluster in similar regions, highlighting the robustness of BayesMonSTR. (**f**) Illustration of two mosaic STR mutations from real single cell data. A 1-bp insertion detected in single-cell colony data is shown on the left. Supporting sequencing reads covering the variant site are displayed for the cell harboring the mutation. A base substitution within STR region, identified from single-cell whole-genome sequencing (scWGS) data is shown on the right. The plots were generated with BayesMonSTR-INSIGHT^9^. (**g**) A gold-standard benchmarking dataset was generated from a breast tissue sample, including tumor epithelial cells, normal epithelial cells and immune cells. Duplicate single-cell libraries, RNA-seq data from the same single cell, and bulk sequencing data from adjacent tissues were generated for cross validation. cross validation, and deep bulk DNA sequencing from adjacent tissues. (**h**) Example variant validated by duplicate library sequencing and RNA-seq from the same cell, with mutant reads also present in adjacent-tissue bulk DNA-seq. Error bars: 95% CI of mutant-cell VAFs (binomial sampling). (**i**) BayesMonSTR achieved multi-fold higher accuracy in detecting cell-specific mosaic STR mutations compared to adapted HipSTR and ExpansionHunter (two-tailed t-test).

For the ground-truth benchmarking, we sequenced 29 single cells from a breast tissue sample, including seven tumor epithelial cells, 12 normal epithelial cells, and ten immune cells (Methods, Supplementary Fig. 2, Supplementary Table 2). Single-cell DNA was amplified using two widely adopted strategies: MDA and PTA (Methods). Across cells, BayesMonSTR consistently achieved ∼60–80% validation rates due to its comprehensive framework—several folds higher than HipSTR(v0.6.2)^6^ and ExpansionHunter(v5.0.0)^8^(p = 8.8 × 10^−9^ for indels and p = 2.1 × 10^−13^ for mismatches, two-tailed t-tests, Fig. 1i, Supplementary Fig. 3), two state-of-the-art germline STR mutation callers that we adapted for mosaic mutation detection by applying a series of hard filters (Methods). Of note, precision and recall were lower in some PTA-amplified cells, likely due to additional PCR cycles (9–10 rounds) that introduced DNA-slippage errors (Supplementary Fig. 4). This superior precision was consistently observed across additional independent benchmarks^10,11^—including public MDA-amplified cells, public single-cell colonies and simulated data—while retaining competitive sensitivity (Methods, Supplementary Tables 3-5, Supplementary Figs. 5-6).

Given the hypermutable nature of STR regions, we further investigated their potential for in vivo lineage tracing. Notably, a phylogeny inferred from shared STR mutations in our benchmark dataset—spanning ∼1% of the genome—roughly recapitulated the topology of a tree constructed from mosaic single-nucleotide variants (SNVs) identified in the remaining 99% of the genome (Methods, Supplementary Fig. 5h, Supplementary Table 6). These results support the use of STR mutations as a cost-effective approach for lineage tracing.

Using this tool, we characterized the landscape of mosaic STR mutations in three mitotic and post-mitotic tissues, including B cells from blood^12^ (56 cells from 14 healthy donors aged 0–106 years), lung epithelial cells^10^ (160 cells from 16 healthy donors aged 1–81 years) and neurons^13^ (126 cells from 19 healthy donors aged 0–104 years) (Supplementary Table 7). The mutation rate per cell per base pair ranges from approximately 10^-6^ to 10^-5^, significantly higher than the reported point mutation rate outside of microsatellite regions^11^. The overall exonic indel mutation rate was about 60% lower than expected (p = 5.5×10^-122^, one-tailed binomial test, Supplementary Table 8). This reduction was not observed for exonic SNVs (p=0.996, one-tailed binomial test), suggesting strong purifying selection specifically targeting exonic STR indels.

Strikingly, neurons—which exit the cell cycle early in development—harbored significantly more mosaic STR mutations than mitotic B cells (p = 2.4 × 10^-6^) and lung epithelium (p = 3.1 × 10^-38^, two-tailed Mann-Whitney U tests, Fig. 2a-b). Size-resolved analysis of indel mutation rates revealed an intriguing dichotomy in age-related dynamics. While B cells (p = 0.003) and lung epithelium (p = 1.8 × 10^-10^) accumulated significantly more 1-bp indels—the most abundant indel type, accounting for 79% of all STR indels—with age (one-tailed t-tests), neuronal 1-bp indel burden showed no age-related increase (p = 0.352, one-tailed t-test, Supplementary Fig. 7a), indicating that 1-bp STR indels were largely acquired during early development in neurons. By contrast, all three tissues, particularly neurons, accumulated longer indels during aging (Fig. 2c, Supplementary Fig. 7b-d). These findings suggest that longer STR indels arise via distinct, age-dependent mechanisms that operate across cell types, whereas the accumulation of 1-bp indels is primarily linked to replicative history. Beyond indels, all three tissues also accumulated significantly more mosaic SNVs with age (Supplementary Fig. 8).

**Fig. 2.**
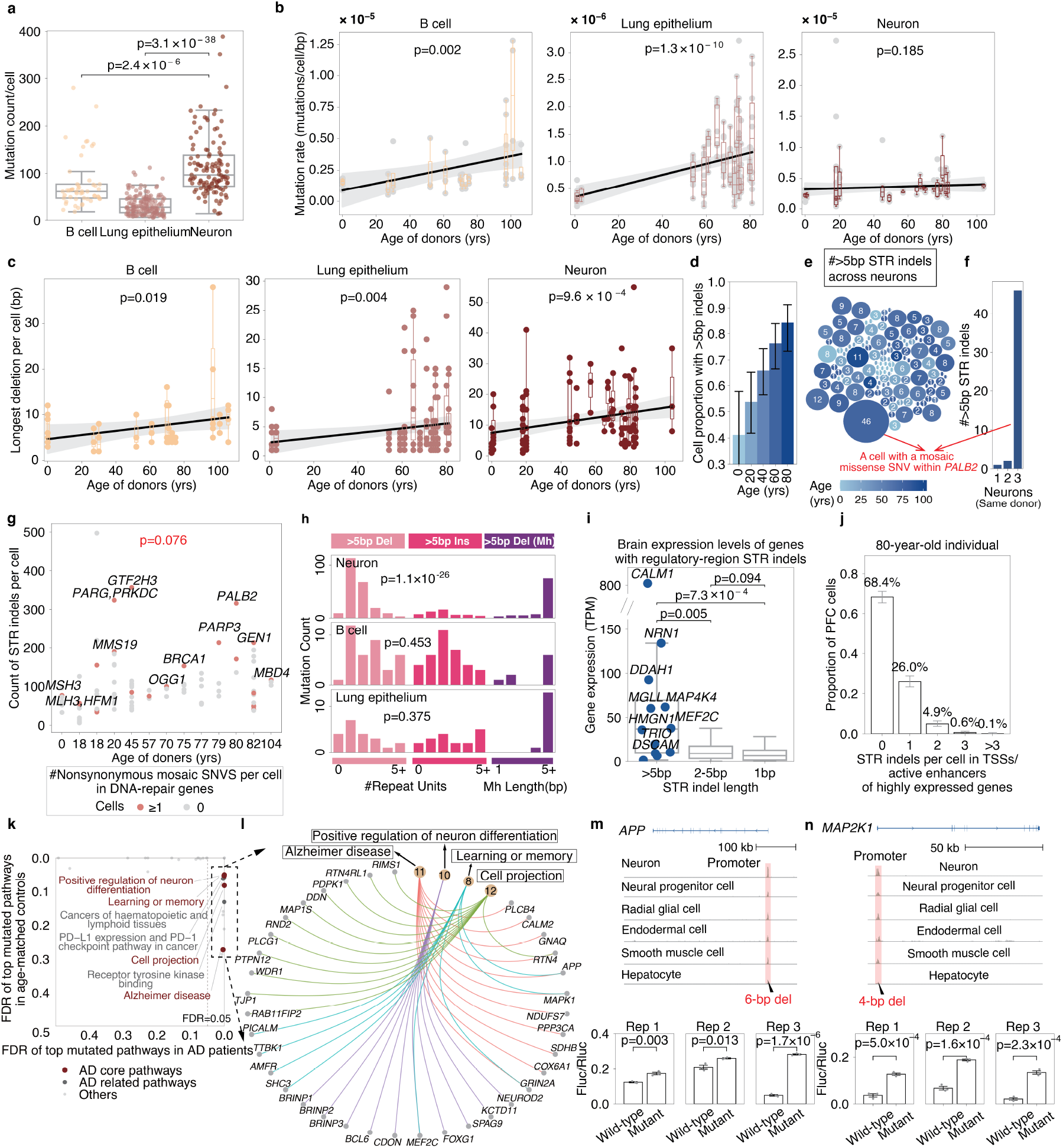
Mosaic STR mutations across tissues in aging and disease. (**a**) Mosaic STR mutation burden (including indels and SNVs) per cell is significantly higher in neurons than in other cell types (two-tailed Mann-Whitney U test). (**b**) Mosaic mutations within and flanking STR regions accumulate with age in B cells and lung epithelial cells but remain stable in non-dividing neurons (one-tailed t-test for slope >0). Shading indicates 95% CIs. Longer STR indels with high signal-to-noise ratios were used in panels c-j and passed manual validation. (**c**) The longest mosaic STR contraction per cell increases significantly with age across all cell types (same regression approach as in b). (**d**) Estimated proportion of neurons carrying mosaic STR indels >5 bp increases with age (binomial Generalized Linear Model (GLM)). Error bars indicate 95% confidence intervals derived from the fitted model. All insertions >5 bp in panels d-g were manually validated. (**e**) STR indels >5 bp are prevalent in neurons, especially with age. Circle size indicates indel count per neuron, color denotes donor age. The neuron with 46 indels carries a missense mutation in DNA repair gene *PALB2*. (**f**) Distribution of >5 bp STR indels in three neurons from the same individual. The *PALB2*-mutant neuron shows substantially higher burden. (**g**) Nonsynonymous mosaic SNVs in DNA repair genes are marginally associated with increased STR indel counts in neurons (Generalized Linear Mixed Models (GLMM) with negative binomial distribution, random intercept for individual). Red points indicate cells with repair-gene SNVs. (**h**) Mutation signatures of >5 bp STR indels across cell types. Mh, microhomology. (**i**) Genes with >5 bp STR indels in TSSs or active enhancers show higher expression than genes with shorter indels (two-tailed Mann-Whitney U test). (**j**) Poisson modeling estimates that >30% of neurons in 80-year-olds carry at least one mosaic STR indel in regulatory regions of highly expressed genes. Error bars indicate 95% confidence intervals computed using binomial Wilson score intervals. (**k**) In AD patients only, STR indels in TSSs and active enhancers are significantly enriched in AD-related genes (Methods). (**l**) Key AD genes contain multi-hit STR indels in regulatory regions across patients. “earlyAD,” “lateAD” indicate disease stage. (**m**,**n**) Luciferase assays in HEK293T cells show that a 6-bp deletion in the *APP* TSS and a 4-bp deletion in the MAP2K1 TSS significantly enhance gene expression (two-tailed t-test, n=3 biological replicates, error bars indicate SD). Normalized H3K4me3 and H3K27ac levels from ENCODE are shown; pink regions indicate promoter/enhancer regions, black triangles indicate indels.

Longer indels affect more genomic sequences and thus pose a greater risk of disrupting DNA structure. Among the three tissues, long STR indels (>5 bp) were significantly more prevalent in neurons than in B cells or lung epithelium (p = 2.8 × 10^-3^ and p = 1.0 × 10^−6^, respectively, two tailed t-tests). By age 80, approximately 84.3% (95% CI: 73.3–91.2%) of neurons from healthy individuals carried at least one such indel (Fig. 2d, Methods). Across 126 neurons from 19 donors, most cells harbored fewer than ten long indels, though a subset showed markedly higher burdens (Fig. 2e). Notably, one neuron from an 80-year-old female exhibited nearly 50 long STR indels—a >30-fold excess over two other neurons from the same individual (Fig. 2f). This same cell also carried a nonsynonymous mosaic exonic SNV in *PALB2* (Supplementary Table 9), a gene essential for DNA repair. More broadly, neurons harboring nonsynonymous SNVs in DNA repair genes showed a marginally significant increase in STR indel burden (p = 0.076, negative binomial GLMM, Fig. 2g, Methods), suggesting that mosaic mutations in DNA repair genes may compromise microsatellite stability during aging.

Notably, among >5bp mosaic indels lacking microhomology at mutation boundaries, >5 bp insertions and deletions occurred at similar frequencies in B cells (p = 0.453) and lung epithelium (p = 0.375), whereas neurons displayed a significant bias toward deletions (p = 1.1 × 10^-26^, two-tailed binomial tests, Fig. 2h), suggesting a specialized long-deletion mechanism in aging neurons. Concurrently, all three tissues accumulated a substantial proportion of >5bp deletions flanked microhomology—a hallmark of microhomology-mediated end joining (MMEJ), which typically generates long deletions. These accounted for 25.9% of all >5 bp indels in neurons, compared to 11.1% in B cells and 26.4% in lung epithelium, suggesting increased double-strand break formation and preferential MMEJ repair in aging tissues^14^.

Beyond indels, mosaic SNVs within STR regions largely mirrored established extra-STR patterns attributed to oxidative damage and cytosine deamination (Supplementary Fig. 9). Together with our indel findings, these results reveal a complex landscape of microsatellite mutagenesis in aging, characterized by both genomic instability and engagement of specific DNA repair processes.

Given the role of microsatellites in gene regulation, we examined STR indel distribution in neuronal regulatory regions. These mutations were enriched in TSSs and active enhancers of genes highly expressed in the brain (p = 5.0 × 10^-3^, one-tailed t-test, Supplementary Fig. 10a), with longer indels showing the strongest enrichment (Fig. 2i). Based on Poisson sampling, over 30% of neurons in an 80-year-old would harbor at least one such indel (Fig. 2j).

To further explore this, we adapted BayesMonSTR for snATAC-seq data (Methods). Analysis of prefrontal cortex from 55 AD patients and 55 controls^15,16^ revealed mutation rates comparable to scWGS data (Supplementary Fig. 10b), with a similar excess of long deletions (p = 1.1 × 10^-31^, one-tailed binomial test, Supplementary Fig. 10c). STR indels in AD patients were significantly enriched in TSSs and active enhancers of AD-related genes and pathways (Fig. 2k-l, Supplementary Fig. 10d), including well-established risk genes such as *APP, CD2AP, PICALM*, and *MEF2C* (Supplementary Table 10). Luciferase assays of six STR indels in TSS/enhancer regions of AD-related genes revealed that four significantly disrupted gene expression in 293T cells (Fig. 2m-n, Supplementary Fig. 11), indicating that microsatellites in active regulatory regions are mutation-prone and can affect expression.

As the first tool for genome-wide detection of somatic STR mutations in single cells, BayesMonSTR enables hypothesis-free discovery of STR mutagenesis in aging and disease. It also provides a unified toolkit for scWGS, snATAC-seq (BayesMonSTR-ATAC), and bulk WGS (BayesMonSTR-BulkMonSTR)^17^, as well as enhanced visualization of STR mutations (BayesMonSTR-INSIGHT)^9^, opening new opportunities to investigate the contribution of STR instability to aging and disease. BayesMonSTR is implemented in Python and R and is licensed under the MIT License. The source code, documentation and examples are available on GitHub at https://github.com/douymLab/BayesMonSTR.

## Data Group

Chengyu Li^6^, Huan Liu^1,2^, Yan Luo^1,2^, Wenlong Li^1,2^, Yangning Lan^1,2^, Xiaodong Liu^1,2^, Shang Cai^1,2^, Danyang He^1,2^, Dan Zhou^9^

9. The Second Affiliated Hospital, School of Public Health, Zhejiang University School of Medicine, Hangzhou, China

## Supplementary Figures

**Supplementary Fig. 1.**
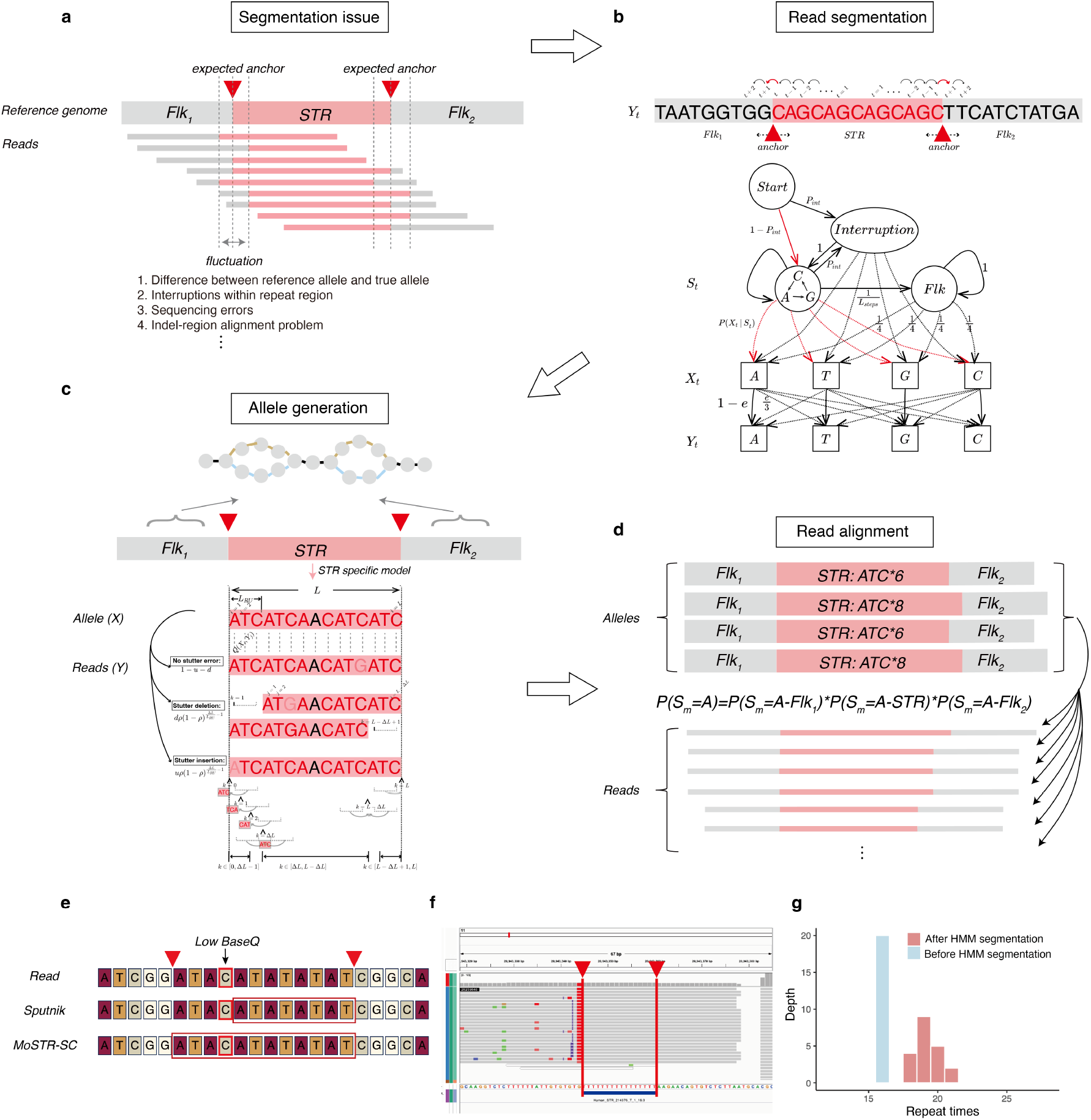
Read segmentation, allele generation and local read alignment based on sequencing reads and information from human reference genome. **(a)** The repeat-flank region boundary (“anchor” in the Figure) separating the STR-region and flanking regions within sequencing reads often fluctuates due to alignment issues, sequencing errors and other problems. **(b)** A fine-tuned HMM was used for reads segmentation. The three hidden states are “STR”, “flanking”, and “interruption”. Each state exhibits a preference for specific base-pair sequences and emits them with a higher probability. The segmentation boundaries between the STR and two flanking regions were identified by enumerating and selecting the optimal hidden state path. **(c)** STR-specialized local read alignment. Assuming no stutter errors occur, the read alignment likelihood was calculated by multiplying the base agreement probabilities between the read sequences and the hidden allele sequences. In the presence of a stutter error, we assume that DNA slippage occurs only once per read and that the stutter-induced INDEL can occur at any position with equal probability. **(d)** We calculate the probability that each read originates from a specific allele using an STR-specialized local read alignment strategy. **(e-g)** The read segmentation strategy of BayesMonSTR is robust to sequencing errors and can accurately restore the correct STR length distribution despite indel fluctuations. **(e)** An example demonstrates how the scoring-based Sputnik algorithm, which ignores base quality, leads to segmentation errors. In contrast, BayesMonSTR’s HMM-based segmentation strategy incorporates base quality, making it more resilient to sequencing errors. **(f-g)** Compared to BWA alignment, which often struggles with accurately defining STR-flank boundaries, the HMM-based approach of BayesMonSTR effectively tolerates indel fluctuations and restores the correct STR length distribution.

**Supplementary Fig. 2.**
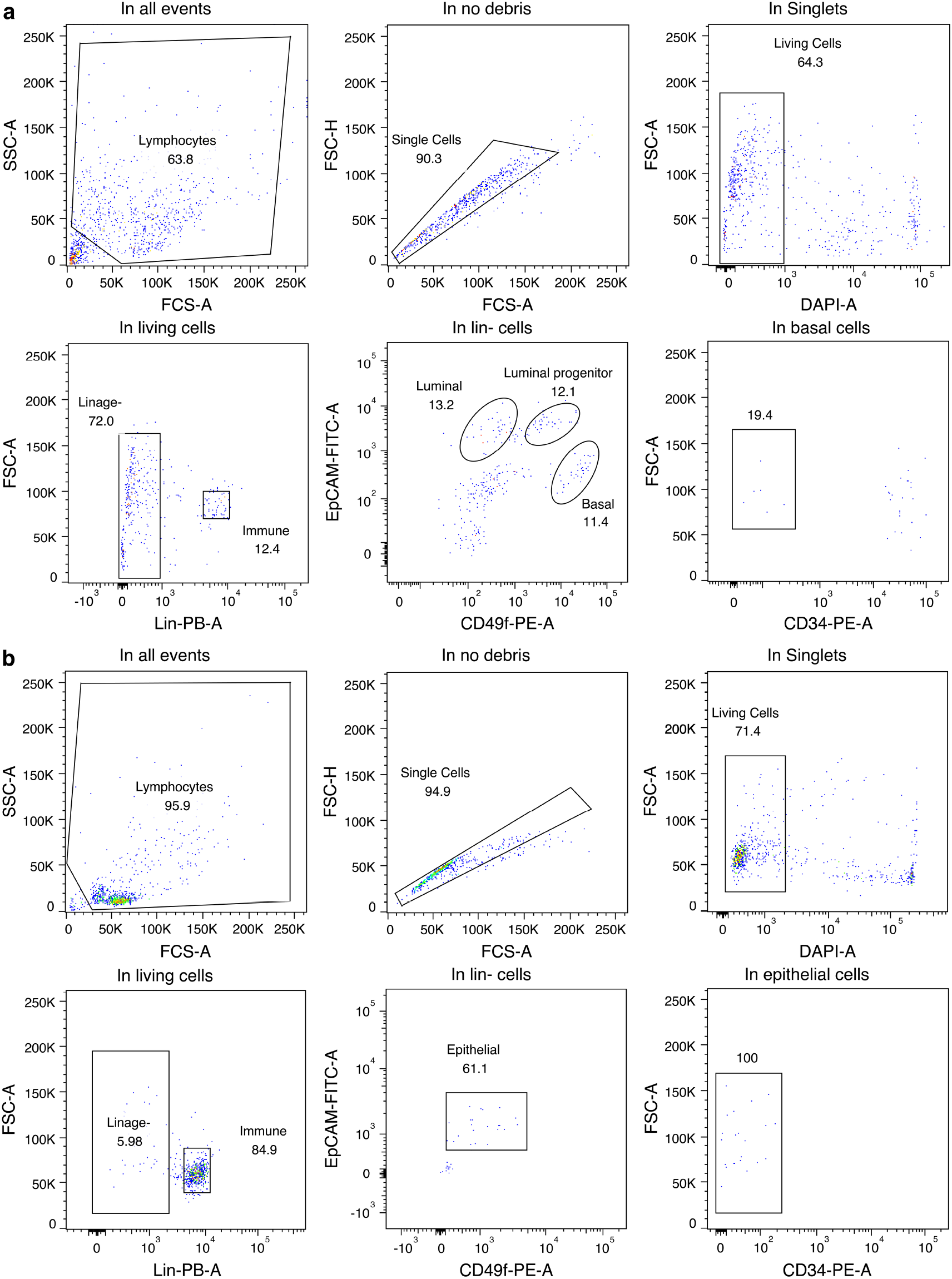
FACS gating strategy for isolating breast-tissue epithelial cells and immune cells. **(a)** Cell sorting results from paired normal tissue samples. Immune cells were defined as Lin+ cell populations, luminal cells were identified as Lin-EpCAM+CD49f-, luminal progenitor cells were identified as Lin-EpCAM+CD49f+, and basal cells were identified as Lin-EpCAM−CD49f+CD34− cells. **(b)** Cell sorting results from tumor samples. Immune cells were defined as Lin+ cell populations, while epithelial cells were identified as Lin-EpCAM+CD49f+CD34-. The sorting strategy was used to isolate these specific cell subsets for further analysis.

**Supplementary Fig. 3.**
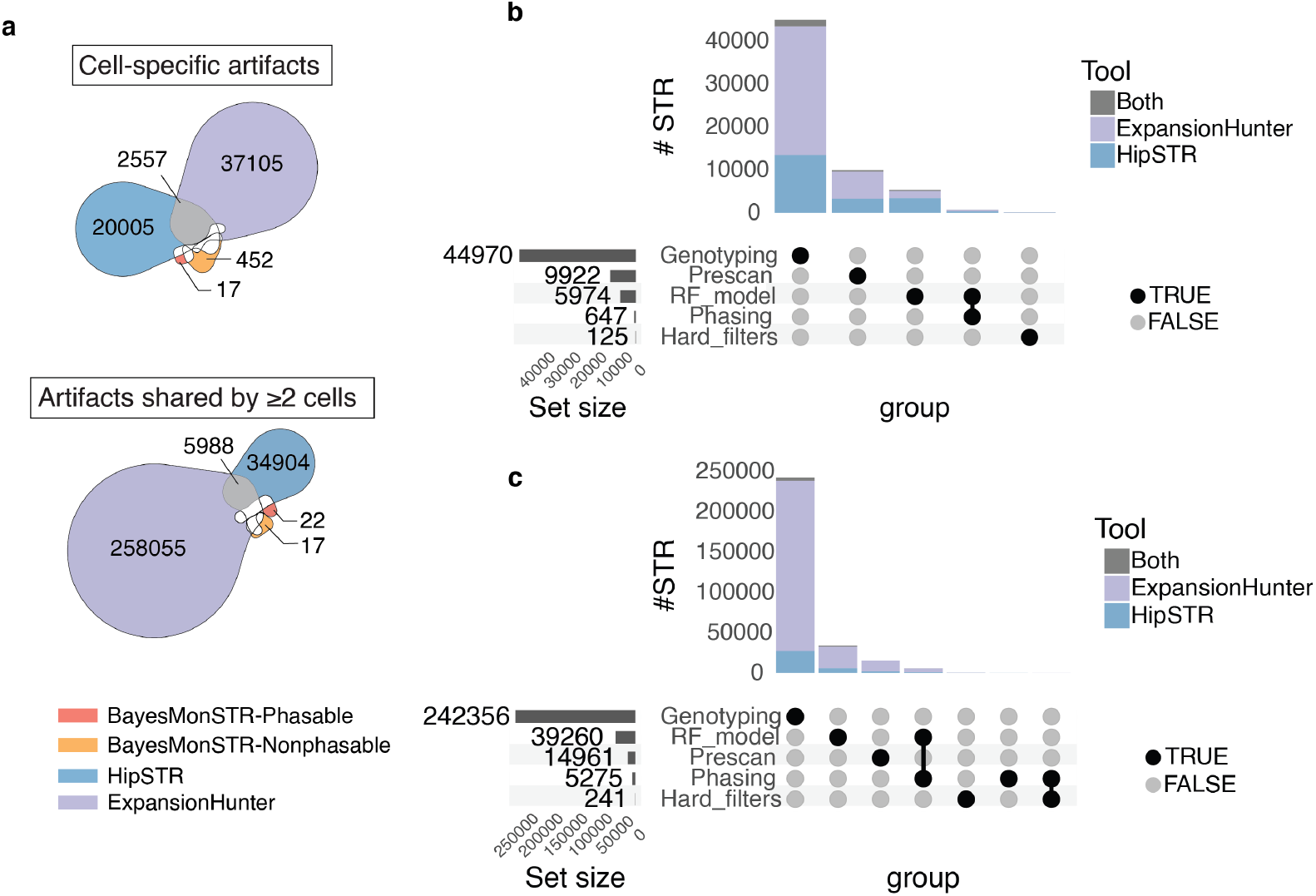
The comprehensive pipeline of BayesMonSTR pipeline effectively excludes artifacts compared with HipSTR and ExpansionHunter. (**a**) Artifact mutations identified by each method for cell-specific (top) and shared (bottom) artifacts from the benchmark dataset we generated. (**b-c**) Artifact mutations from HipSTR and ExpansionHunter were not called by BayesMonSTR through the multiple steps (Genotyping, RF model, Prescan, Phasing, Hard_filters) for cell-specific (**b**) and shared (**c**) artifacts.

**Supplementary Fig. 4.**
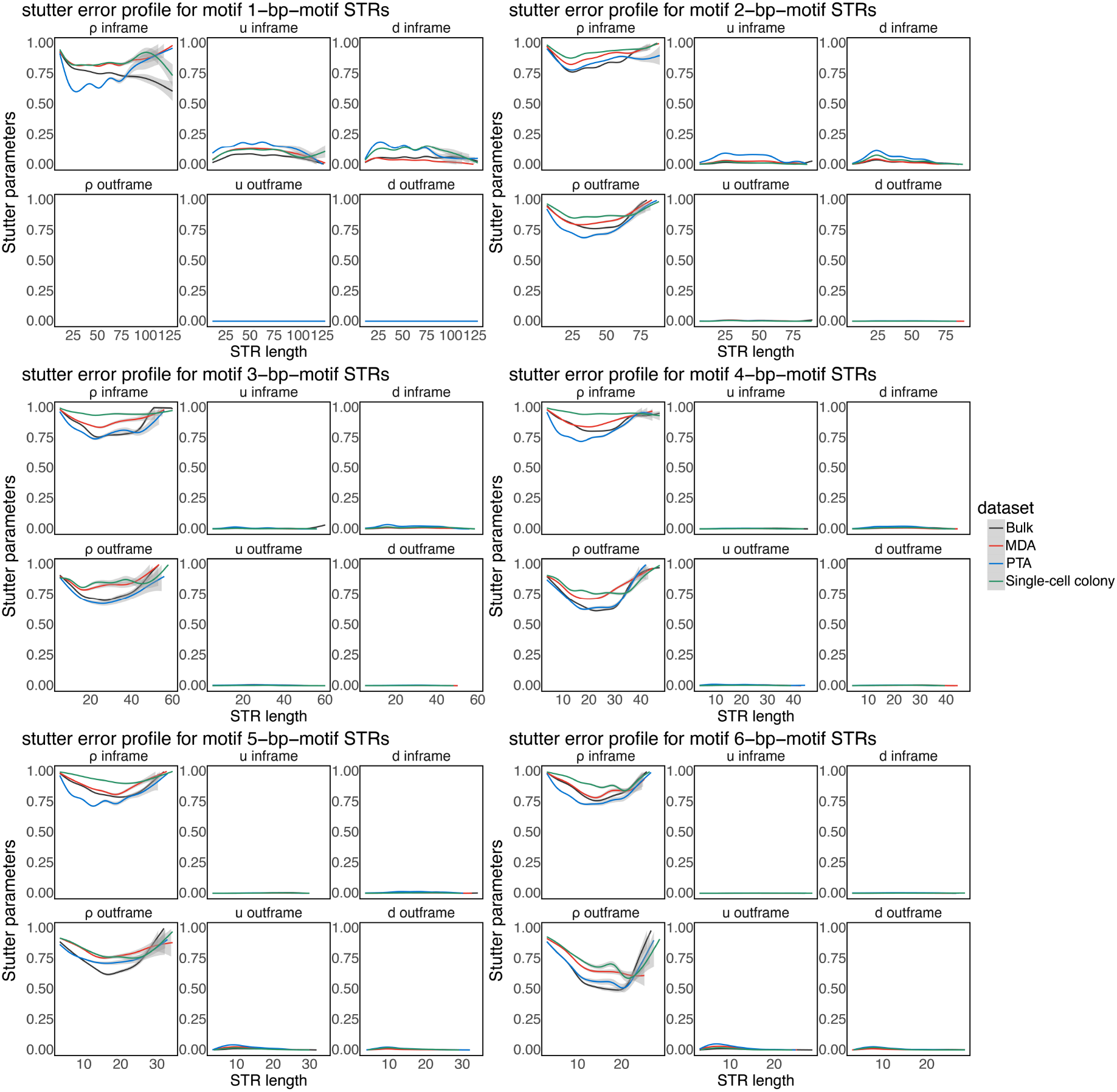
Stutter error estimation across bulk DNA sequencing data, MDA cells, PTA cells and single cell colonies. Stutter-error parameters were derived from a length-based stutter error geometric distribution model, calculated with an EM-algorithm. “Bulk” refers to bulk DNA-seq data, “MDA” refers to single-cell sequencing data amplified with multiple displacement amplification approach, “PTA” refers to single-cell sequencing data amplified with primary template-directed amplification approach, and “SCC” refers to single-cell colony data. STR loci were described by the parameters *ρ* (rho), *u* (up), and *d* (down), which represent the three parameters of the geometric distribution, in-frame and out-frame parameters were estimated separately (Methods). Smoothed trend lines were calculated by the generalized additive model (GAM) with cubic splines, and gray area represents 95% confidence interval.

**Supplementary Fig. 5.**
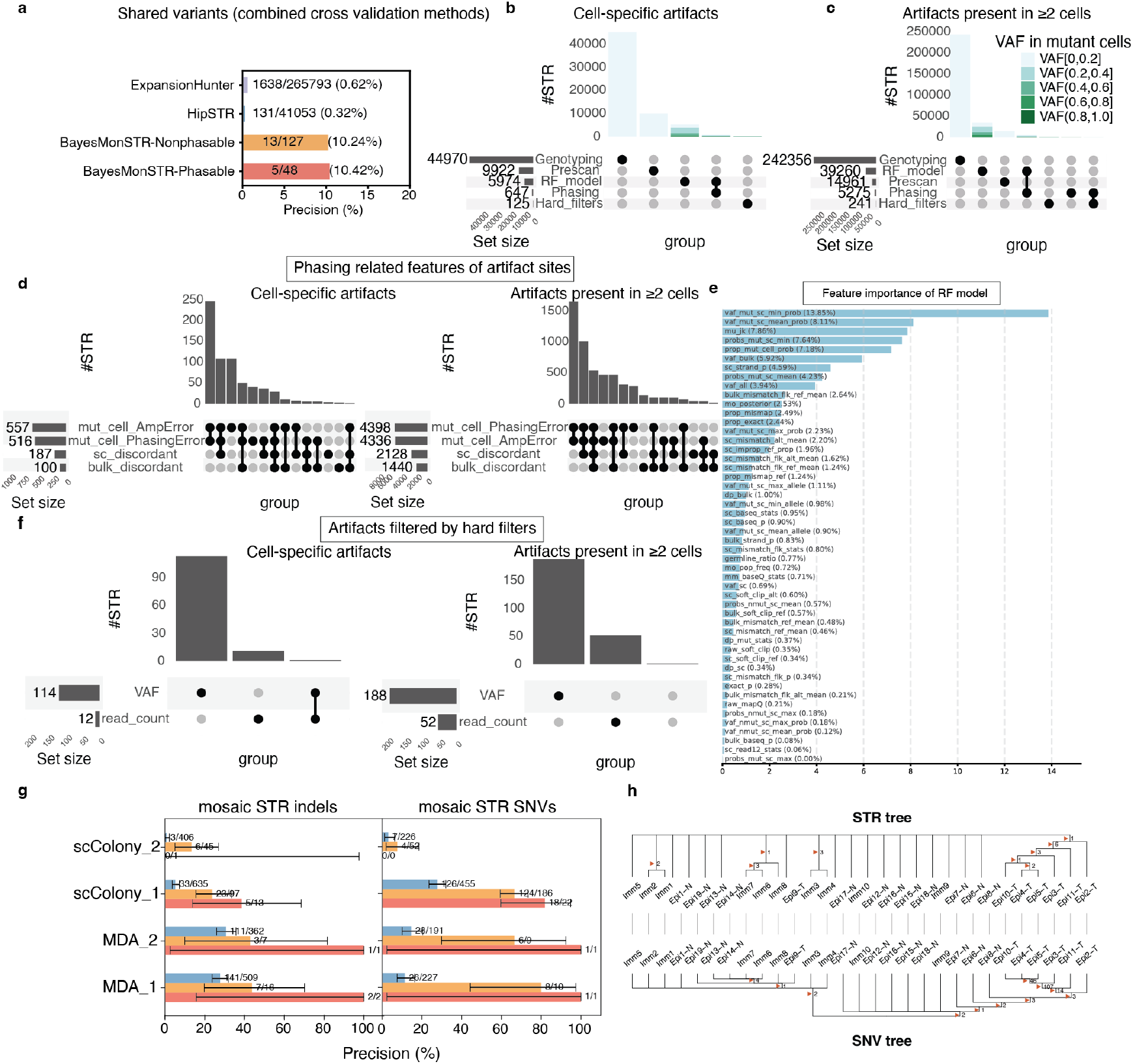
BayesMonSTR outperforms HipSTR and Expansionhunter in detecting mosaic STR mutations from single cells. **(a)** Validation rate of mutations presents in ≥2 cells across different software tools. The sites were extensively evaluated using both orthogonal data and checked in the phylogeny tree reconstructed with PhyloSOLID (Methods). **(b-c)** Distribution of mutant-cell VAFs for false-positive sites detected by HipSTR and ExpansionHunter. Solid points indicate false positives that were filtered by BayesMonSTR using a specific filtration method. It is evident that BayesMonSTR excluded a majority of these false positives, which had relatively low mutant allele frequencies, during the pre-scan and genotyping stages. Mutations with relatively high mutant-cell VAFs were further excluded by BayesMonSTR using the RF classification model, probabilistic haplotype phasing, and a series of hard filters. **(d)** Expanded analysis of false-positive sites identified by HipSTR and ExpansionHunter excluded by phasing-related features of BayesMonSTR. Features driving the exclusion of false positives by include discordant read pair proportion, amplification error-related features. (**e**) Key features of the random forest model of BayesMonSTR. False positives were effectively filtered by BayesMonSTR by using a combination of informative predictors. Explanations of these predictors are provided in Supplementary Table 1. **(f)** Artifact sites excluded by BayesMonSTR using hard filters. **(g)** Performance benchmarking of BayesMonSTR versus HipSTR using additional public datasets. Error bars represent 95% confidence intervals of the validation rates calculated using binomial sampling. MDA_1 (UMB1465_21-ctx-1cP2B11_1946-WGS-PFC_19), MDA_2 (UMB1465_21-ctx-1cP2F06_1946-WGS-PFC_20), scColony_1 (10_PRL3-1_001B8_sc16), and scColony_2 (10_PRL3-2_002E11_sc17). (**h**) A phylogeny tree reconstructed using shared mutations from our benchmark shows that the mosaic STR tree roughly resembles the mosaic SNV tree. Shared STR variants were identified using only the 16 MDA cells, due to the substantially higher PCR stutter error rate inherent to PTA-amplified cells.

**Supplementary Fig. 6.**
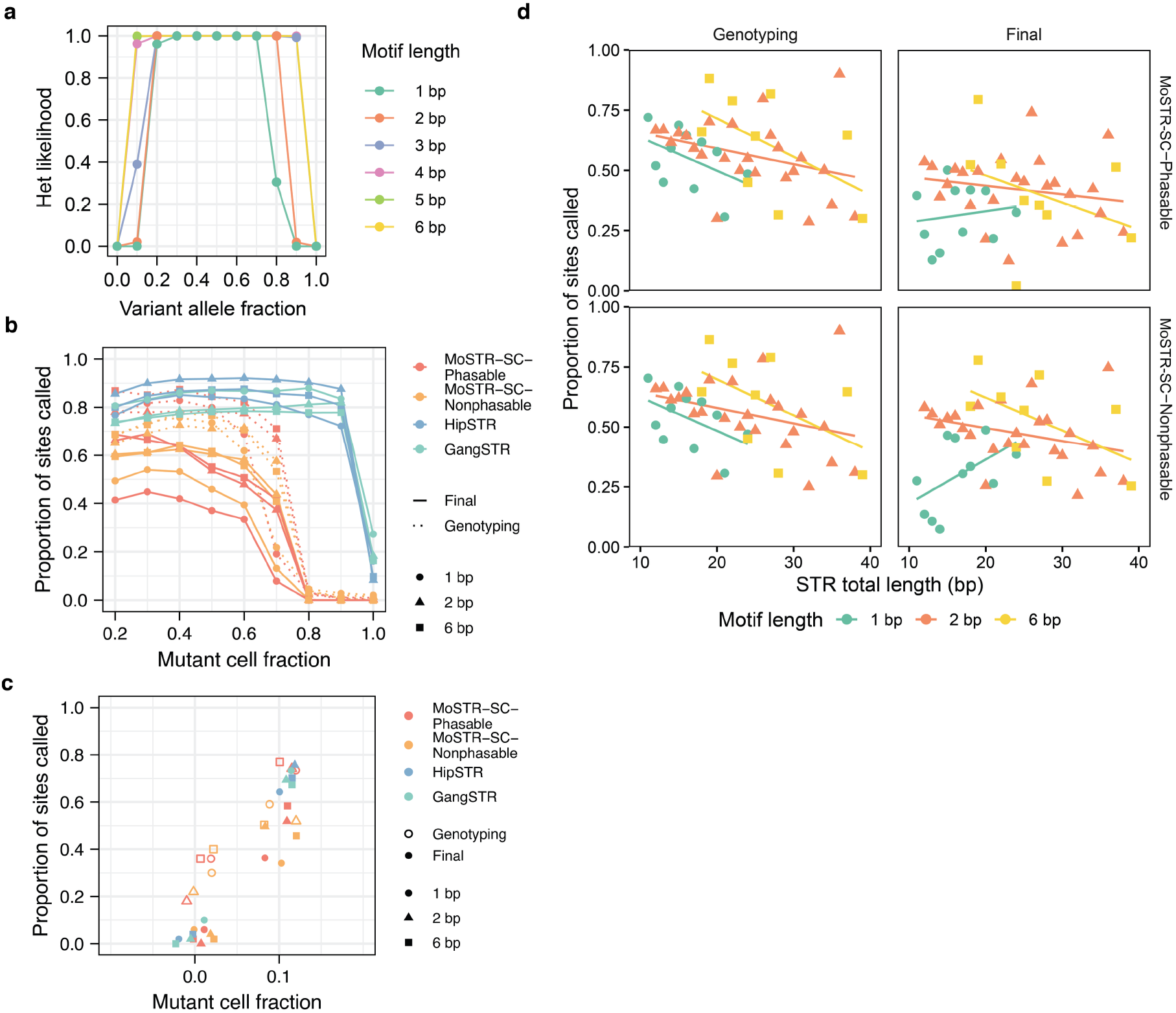
Performance evaluation of BayesMonSTR using simulated data with spike-in mutations. **(a)** Genotyping likelihoods of germline heterozygous at different variant allele fractions. Reads are assumed with Phred quality score 30 and median stutter error parameters were used for likelihoods calculation, heterozygous likelihoods were normalized along with reference and alternative homozygous likelihoods. **(b-c)** Sensitivity comparison across the two software tools. HipSTR was chosen for the benchmark here, given its superior performance to ExpansionHunter on the real-world data. One-motif mosaic STR insertions within 16-24 bp STR regions were generated using BAMSurgeon. The total number of cells was 10. Dashed lines indicate the raw mutation callings after genotyping of BayesMonSTR, while solid lines indicate final call sets. B illustrates the sensitivity detecting shared mutations and C illustrate cell-specific mutations. **(d)** Sensitivity variation across different STR total lengths. The total cell number was 10 and mutations spiked in were 1-motif insertions.

**Supplementary Fig. 7.**
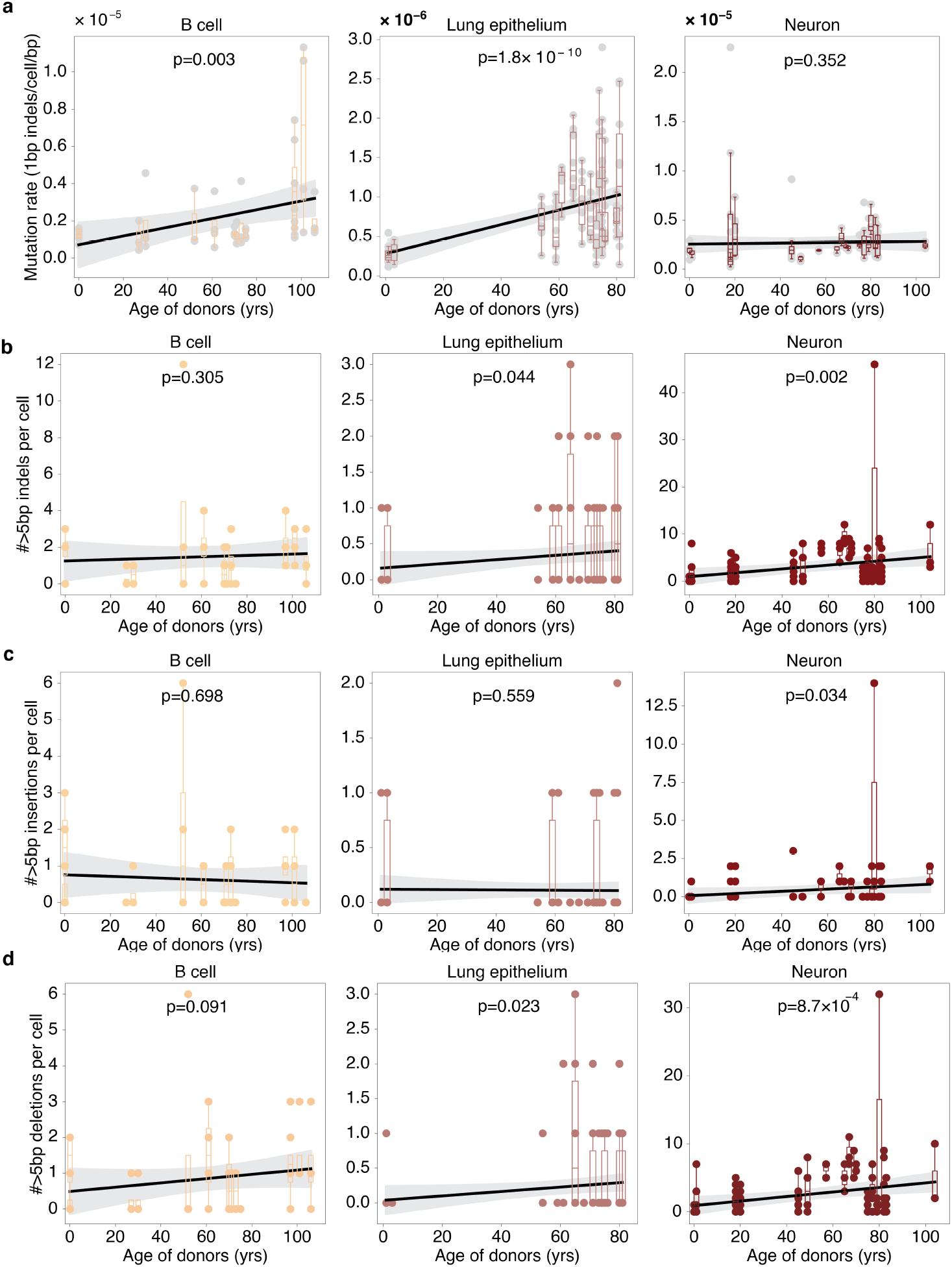
Mosaic STR indels across cell types. **(a)** 1-bp STR indels accumulate with age in B cells and lung epithelial cells. In contrast, the mutation rate remains relatively stable in non-dividing neurons across donors of different ages. **(b–d)** Relationship between age and the total burden per cell of STR indels >5 bp (b), STR insertions >5 bp (c), and STR deletions >5 bp (d) across three cell types. In all three cell types, the burden of STR deletions >5 bp shows a positive correlation with age. However, only in neurons does the burden of indels, insertions, and deletions consistently increase with age. Linear regression equations were derived using the least squares method. P-values were calculated using a one-tailed t-test to test whether the regression coefficient is significantly greater than zero. The shaded areas around each curve represent the 95% CIs, assuming the differences between predicted values and observed values follow a t-distribution.

**Supplementary Fig. 8.**
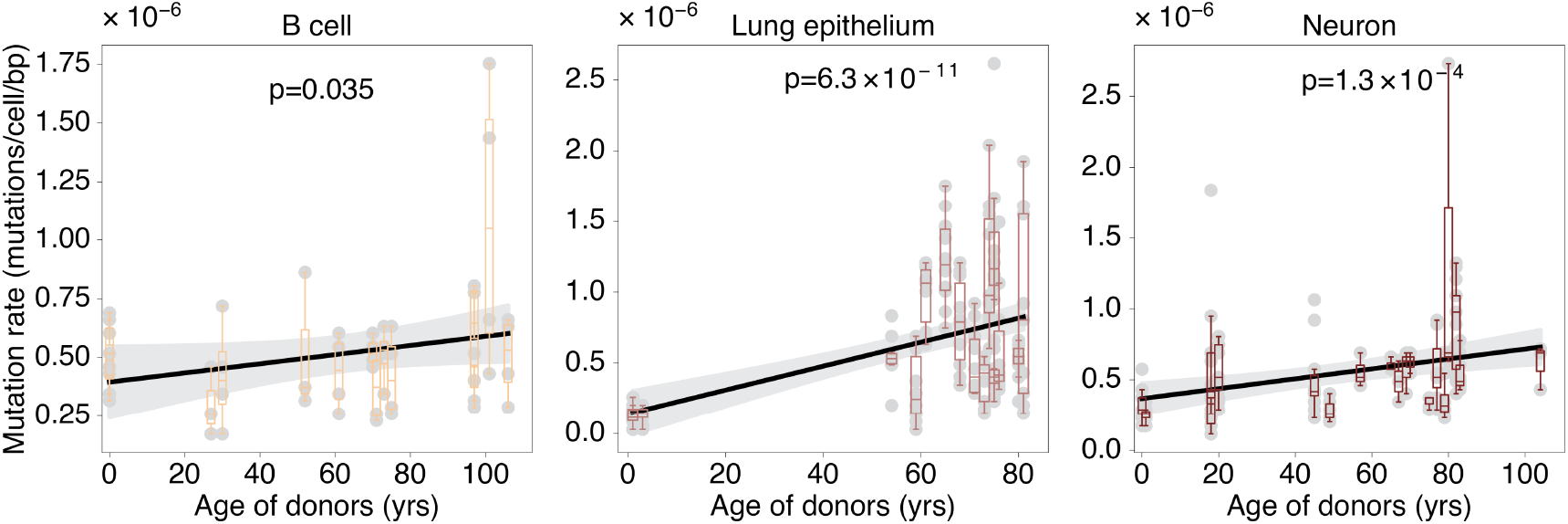
Mosaic STR SNVs across cell types. STR SNVs accumulate with age in all three cell types, as illustrated using the methodology described for Supplementary Figure 7.

**Supplementary Fig. 9.**
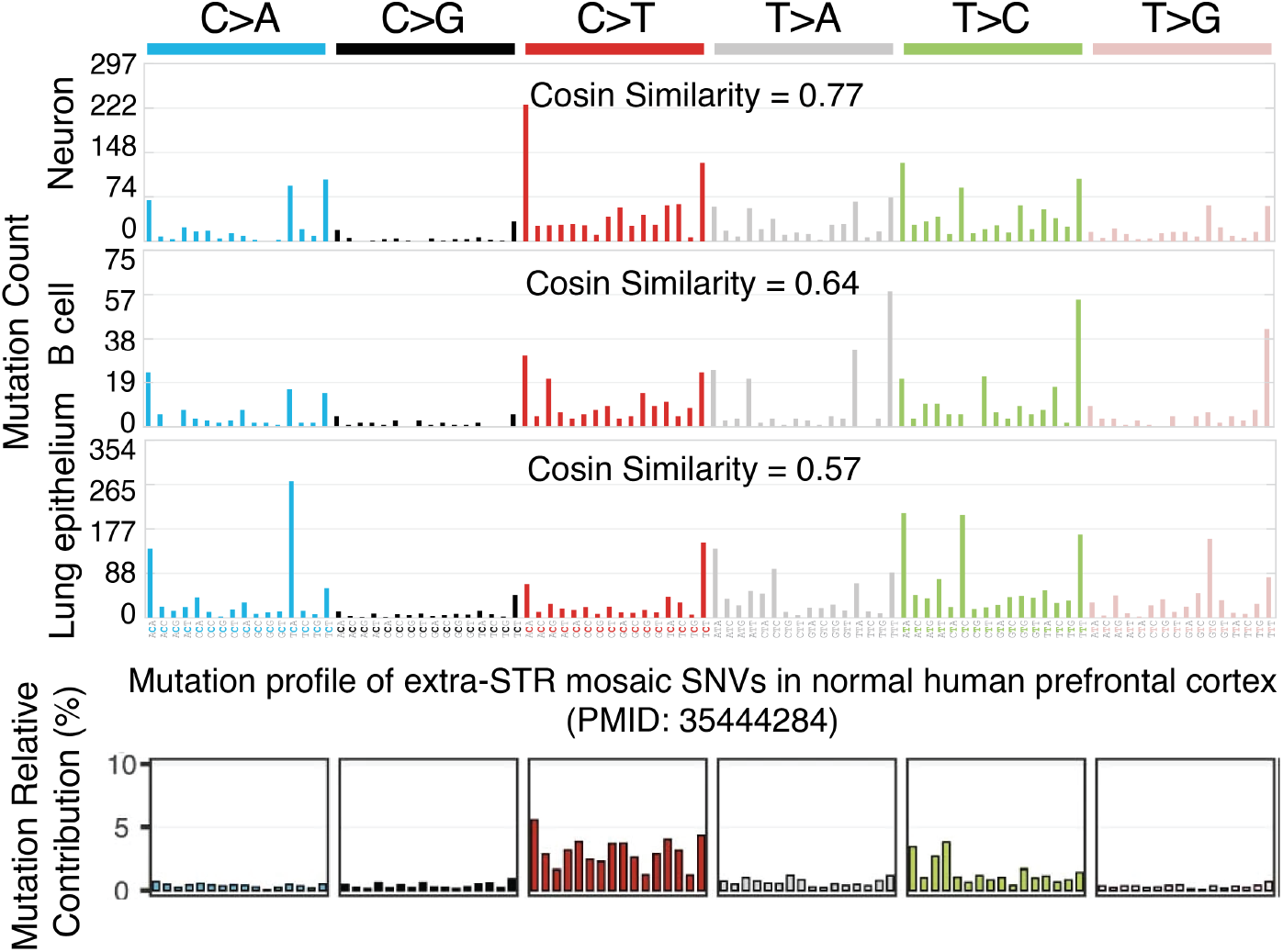
Mutational mechanism analysis of mosaic STR mutations. STR-region SNVs largely mirrored the established patterns of extra-STR mosaic SNVs in normal human prefrontal cortex from PMID 35444284. The mutation signature similarity was computed via cosine similarity.

**Supplementary Fig. 10.**
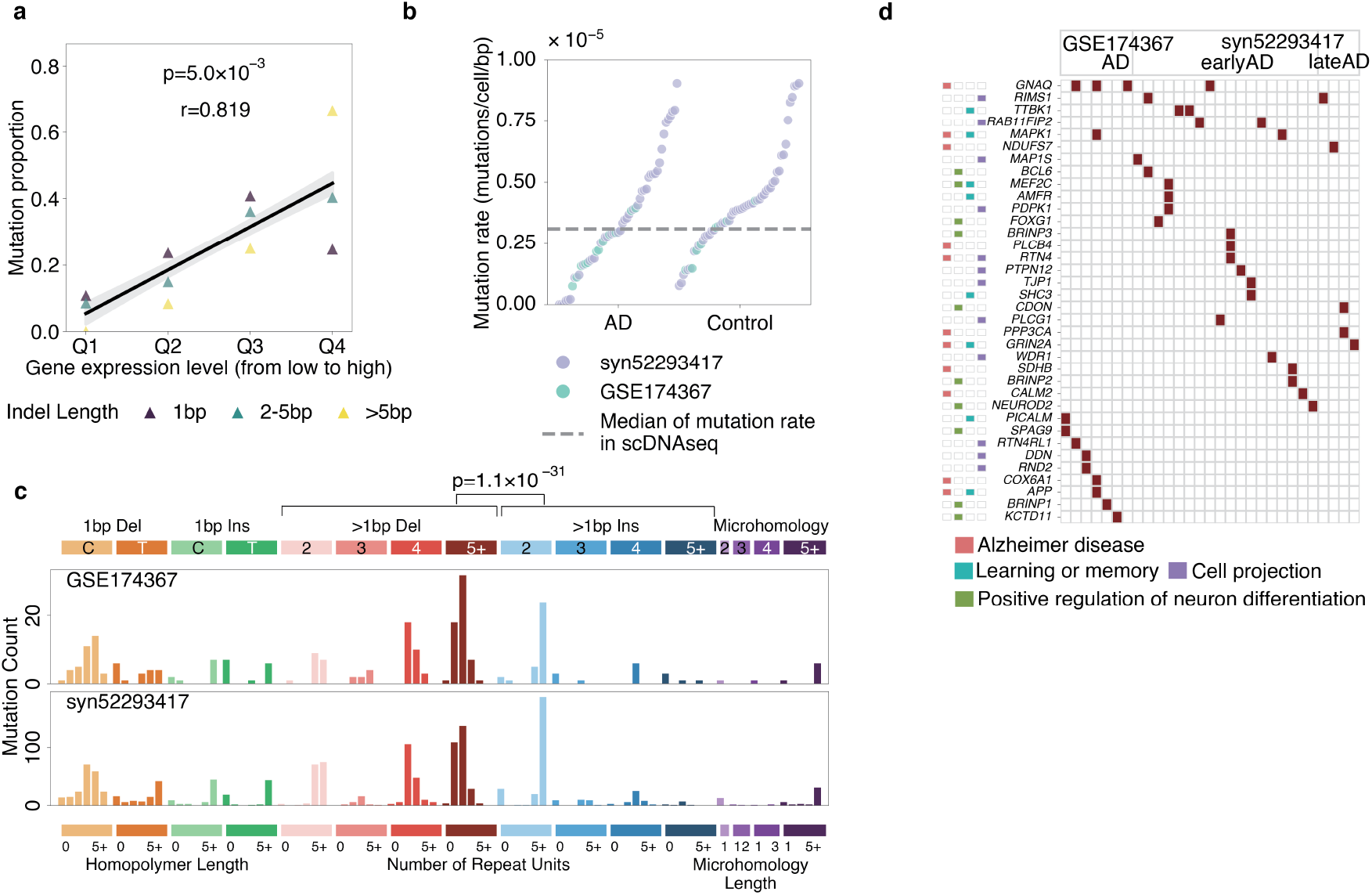
Regulatory mosaic STR indels in neurons. **(a)** Mosaic STR indels in neurons are enriched in TSSs and active enhancers of highly expressed genes. The number of variants is indicated beside each data point. Q1-Q4 represent expression level quantiles from low to high (Methods), and the brain gene expression data were obtained from ENCODE. Linear regression equations were derived using the least squares method. P-values were calculated using a one-tailed t-test to test whether the regression coefficient is significantly greater than zero. The shaded areas around each curve represent the 95% CIs, assuming the differences between predicted values and observed values follow a t-distribution. **(b)** snATAC-seq samples exhibit mutation rates highly consistent with those observed in scWGS data. **(c)** A significant excess of >1 bp deletions over >1 bp insertions was observed in snATAC-seq data. P-value was calculated using a one-tailed Binomial test. **(d)** Multiple key genes in AD pathways contained multi-hit STR indels in their active regulatory regions across different patients. “earlyAD” and “lateAD” indicate early-stage AD patients and late-stage AD patients.

**Supplementary Fig. 11.**
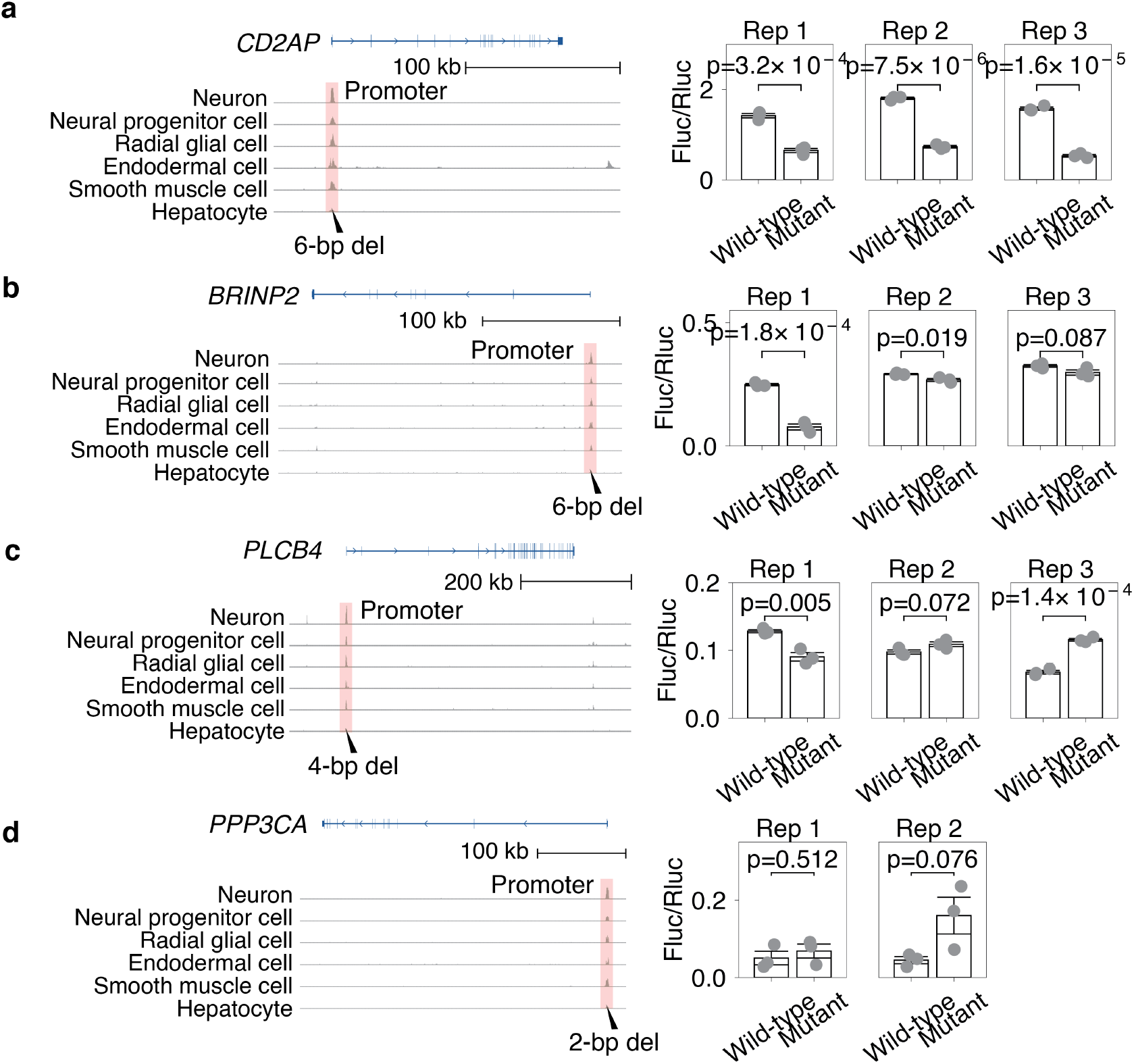
Luciferase assay results of somatic STR indels in regulatory regions. Normalized levels of H3K4me3 and H3K27ac—key histone modifications linked to promoter and enhancer activities across cell types were obtained from the ENCODE project. Pink regions indicate the promoter or enhancer regions of the gene, with indels highlighted by black triangles. Luciferase assays in HEK293T cells revealed that a 6-bp deletion in the TSS of *CD2AP* and a 6-bp deletion in the TSS of *BRINP2* significantly disrupted gene expression (P < 0.05, two-tailed t-test). Furthermore, a 4-bp deletion in the TSS of *PLCB4* and a 2-bp deletion in the TSS of *PPP3CA* enhanced gene expression with marginal significance (P < 0.1, two-tailed t-test). Error bars represent standard deviation (3 biological replicates).

## Methods

### Sequence alignment and data preprocessing

Raw FASTQ reads were aligned to the GRCh37 reference genome (human_g1k_v37_decoy) using BWA-MEM (v0.7.18). Duplicate reads were marked with Picard MarkDuplicates (v2.27), and base quality score recalibration (BQSR) was performed with the Genome Analysis Toolkit (GATK v4.2). Germline variants were joint called from MDA cells and matched bulk sequencing data with GATK HaplotypeCaller and filtered using Variant Quality Score Recalibration (VQSR) following GATK Best Practices. Common germline heterozygous SNPs (gnomAD^1^ allele frequency >0.1%) were selected for downstream haplotype phasing with Eagle2 (v2.4.1)^2^.

### Stutter-error estimation

To reduce amplification- and sequencing-related artifacts introduced by *in vitro* polymerase slippage, BayesMonSTR models STR stutter errors using a geometric-distribution^3^. The stutter error probability *p* (*r*_*m,c*_ | *a* = *j*) for *m*-th read *r* for cell *c* in the given allele *j* was modeled as follows:

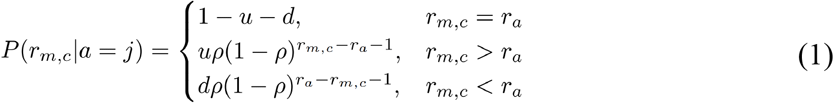

Where:

○ *u* denotes the total mutation rate of artifact insertions.
○ *d* denotes the total mutation rate of artifact deletions.
○ *ρ* denotes the success probability of the geometric model, which belongs to the interval (0,1].

Assuming independence across reads, the genotyping likelihood for each diploid cell is computed using equation 2:

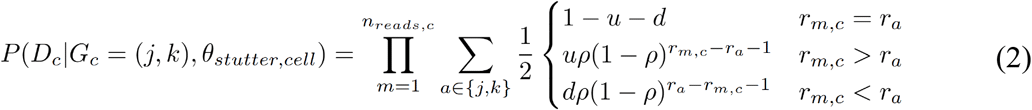

Where:

○ *G*_*c*_ denotes the genotype of a given cell.
○ *j* and *k* denote the two haplotype alleles.
○ *D*_*c*_ denotes the sequencing reads of a given cell *c*.
○ The parameter set *θ* denotes the stutter model parameters (*u, d, ρ*), as described in formular (1).

Of note, reads with mapping quality < 20, as well as reads with mean base quality < 20, were excluded before stutter-parameter estimation.

#### Parameter initialization

○ Given *n* candidate alleles, the initial population frequency for allele *a, f*_*a*_, was set to 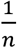, representing equal prior probability across all alleles.
○ The initial mosaic allele fraction parameter µ_msi_ was set to 0.001.
○ Stutter error parameters *u, d* and *ρ*, which define the geometric model, were initialized to 0.01, 0.01, and 0.9, respectively.

#### E-step

In the E-step, posterior probabilities of individual germline genotypes were computed using equation 3:

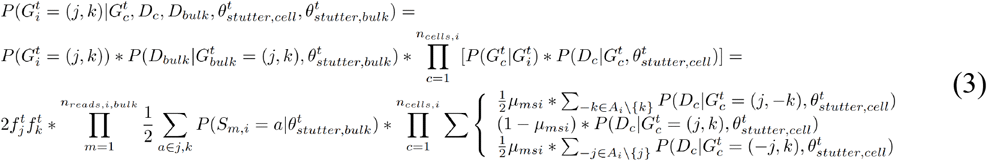

Where:

○ *f*_*j*_ denotes population allele frequency for allele *j*.
○ *G*_*i*_ denotes the individual germline genotype for individual *i*, and *G*_*c*_ denotes the cell genotype.
○ *θ* denote the set of stutter error parameters as previously described.
○ *D*_*c*_ denotes the sequencing reads of a given cell *c*.

Here we assume *G*_*i*_ is the same as *G*_*bulk*_, and the prior probabilities of individual genotypes were calculated based on Hardy-Weinberg Theorem. When *j* equals to *k*, the prior probability of a homozygous germline genotype is given by 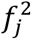.

#### M-step

We then estimated the error parameters *u, d* and *ρ*, and the length-based population allele frequency *f* at *t* + 1-th step for stutter errors in the M-step, and were calculated by equations 4-7, as follows:

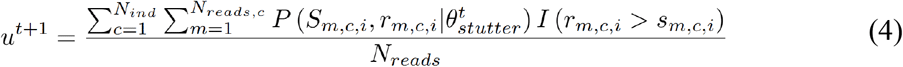

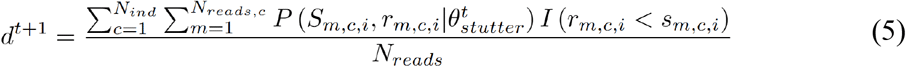

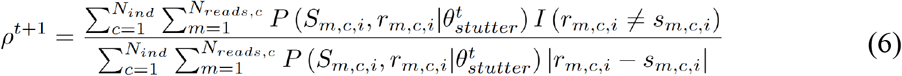

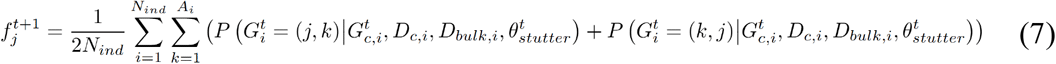

Where:

○ *N*_*ind*_ denotes total number of individuals.
○ The indicator function *I* is used to restrict the *c*omputation to reads with insertion, deletion and nonzero stutter for *u, d* and *ρ* respectively.
○ *p*(*S, r*|*θ*) is read assignment probability for read *r*_*m,c*_ given allele pool *A*_*i*_ (see formula 8).
○ *G*_*i*_ denotes the individual germline genotype for individual *i*, and *G*_*c*_ denotes the *c*ell genotype.
○ *θ* denote the set of stutter error parameters as previously described.
○ *D*_*c*_ denotes the sequencing reads of a given *c*ell *c*.
○ *N*_*reads*_ denotes the total number of samples with the same sequencing method. (including bulk, WGA, single *c*ell *c*olonies, etc).
○ *N*_*reads,c*_ denotes the total number of reads for sample *c*.
○ the indicator function *I* is used to restrict the computation to reads with insertion.
○ deletion and nonzero stutter for *u, d* and *ρ* respectively.
○ *θ* denoted these stutter parameters.
○ *f*_*j*_ denotes population allele frequency for allele *j*.

*p*(*S, r*|*θ*) is read assignment probability for read *r*_*m,c*_ given allele pool *A*_*i*_:

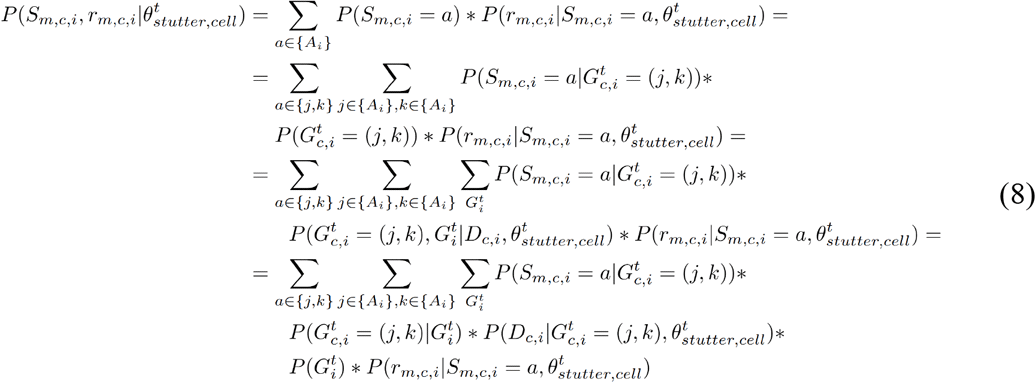

Where:

○ *a* is one of the alleles in the alleles pool *A*_*i*_ = {j, *k*} for individual *i*.
○ other parameters are defined in Equation (3)

### Probabilistic read segmentation

Accurate detection of repeat regions within sequencing reads is essential for generating candidate STR alleles. Traditional segmentation methods, which rely on alignment coordinates and repeat patterns, often suffer from inaccuracies due to mapping biases and interruptions. These issues lead to fluctuations in the ‘anchor points’ used to distinguish repeat and flanking regions, causing the anchor points to deviate from their expected coordinate across different reads (Extended Data Fig. 1a). To address these issues, we utilized a Hidden Markov Model (HMM) approach to identify repeat regions in sequencing reads.

Notably, our HMM model integrates information from both sequencing reads (including sequencing data and quality scores) and the reference genome (repeat-region and flanking region sequences). It also incorporates prior knowledge of in-frame and out-of-frame interruptions derived from the human reference genome. This design enables robustness to sequence interruptions, sequencing errors, and mutations within STR regions, while integrating base quality scores directly into the segmentation process (Extended Data Fig. 1).

Specifically, the HMMs were designed to model the STR sequences using three hidden blocks (Extended Data Fig. 1b):

A. STR block: Represents perfect repeat motifs, capturing the repetitive structure at each STR locus. For example, the motif CAG is modeled with three hidden states— S_C_, S_A_, and S_G_ —within this block.
B. Interruption block: Models mismatches and indels within repeat regions. Similar to profile HMM^4^, this block accounts for variations inside the repeat tract.
C. Flanking block: Represents sequences flanking the STR. In addition to a general flanking state with equal emission probabilities for A, C, G and T, we introduce a locus-specific 3-bp known flanking states extracted from the human reference genome. These states incorporate sequence context to improve segmentation accuracy.

To initialize the transition probabilities, the initial transition probability from the STR block to the flanking block was defined as 1/L, where L corresponds to the expected segmentation distance before entering the flanking region. For homopolymers, L was set to 5 bp, while for non-homopolymers, L was set to twice the motif length. Furthermore, the initial transition probability from the STR block to the interruption block is estimated through an extensive grid search using a reference panel^3^.

The emission probability matrix integrates both the expected STR motif sequence and base quality scores to improve robustness against sequencing errors. Each hidden state corresponds to an expected nucleotide that receives the highest emission probability. In the flanking block, known 3-bp flanking sequences from the reference genome are assigned predefined emission probabilities, while nucleotides outside this region have equal probabilities across A, C, G, and T. For a given hidden state, if the expected base is *X*_*t*_, the observed base is *Y*_*t*_, and the sequencing error rate is e, the emission probability for this base is calculated as follows:

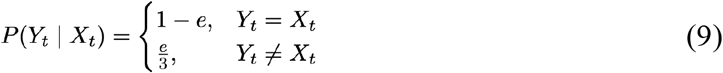

Where:

○ *Y*_*t*_ indicates the observed base in sequencing reads.
○ *X*_*t*_ indicates the expected base given a specific hidden state.
○ *e* indicates sequencing error rate.

Finally, the boundaries between STR and flanking regions are determined by finding the maximum-likelihood hidden state path using the Viterbi algorithm. Based on this optimal state path, sequencing reads are segmented into STR and flanking segments. In cases of large deletions where the mutant allele length falls below the predefined step length—leading to potential segmentation failure—a BWA-based coordinate mapping approach is used as a backup. Similarly, due to the high sequence complexity at imperfect STR loci with 4-bp, 5-bp, or 6-bp motifs containing interruptions, segmentation performance may be compromised; for such loci, the BWA-based coordinate method is also applied.

### STR-specific read alignment

STR sequences from the segmented reads are used to generate candidate alleles, each comprising the core STR region and 5 bp of flanking sequence on both sides. Segmented reads are subsequently aligned to the candidate alleles using a STR-specific alignment method^3^ to calculate the alignment likelihood. This method accounts for both potential PCR stutter errors and sequencing errors. Specifically, three possible scenarios are considered for the sequencing reads: no stutter error, stutter deletion, and stutter insertion. The calculation of alignment likelihood varies slightly for each scenario, as illustrated in Extended Data Fig. 1c.

Given a haplotype *X* of length *L*, if read *Y* is assumed to be free of stutter errors, the likelihood of the observed alignment is determined by the agreement between each base i in the read and its corresponding base in the haplotype:

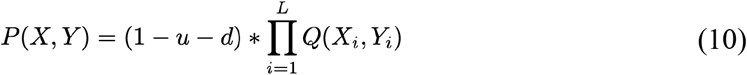

Where:

○ *X* denotes the given STR haplotype.
○ *Y* denotes the STR haplotype observed in the read.
○ *L* denotes the STR length of *X* haplotype.
○ *u* denotes stutter insertion rate from the estimated stutter model.
○ *d* denotes stutter deletion rate from the estimated stutter model.
○ *Q*(*X*_*i*_, *Y*_*i*_) represents the agreement score between the *i* -th base of haplotype *X* and the *i*-th base of haplotype *Y*,as defined in Equation (9).

Otherwise, if read *Y* contains a stutter deletion of haplotype *X*—with a deletion size of Δ*L*—we assume that the deletion could occur at any position k within the haplotype, where k ranges from 1 to total_haplotype_length - Δ*L* + 1 . The read alignment likelihood is then determined jointly by the probability of stutter deletion and the base-wise agreement between the read and the haplotype. The overall likelihood is obtained by averaging over all possible deletion positions:

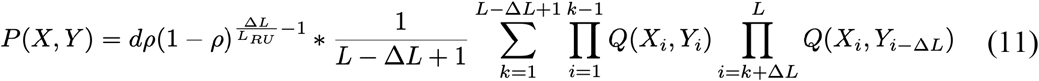

Where:

○ Δ*L* denotes deletion size in read *Y*.
○ *L*_*RU*_ denotes the motif length (in bp).
○ *ρ* denotes the success probability of the geometric model, which belongs to the interval (0,1].
○ *k* denotes the deletion position in the haplotype *X*.
○ Other parameters are defined in Equation (10).

In the third scenario, where the read *Y* results from a stutter insertion in haplotype *X* (insertion size Δ*L*), we assume the inserted sequence arises from a local duplication of adjacent sequence. Consequently, the duplication can originate from either the left or right flank of the insertion site. Due to boundary constraints—insertions within the first Δ*L* positions can only be duplicated from the right flank, and those within the last Δ*L* positions only from the left—only one duplication direction is considered for such edge cases. The alignment likelihood is determined jointly by the stutter insertion rate and the base-wise agreement between the read and the haplotype. As before, the overall likelihood is obtained by averaging over all possible insertion positions and duplication directions:

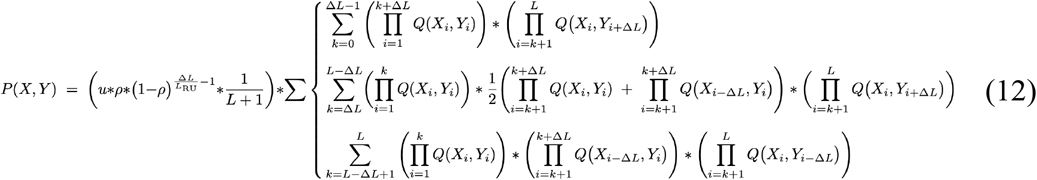

Where:

○ Δ*L* denotes the insertion size in read *Y*.
○ *L*_*RU*_ denotes the motif length.
○ *ρ* denotes the success probability of the geometric model, which belongs to the interval (0,1].
○ *k* denotes the insertion position in the haplotype *X*.
○ Other parameters are defined in Equation (10).

We use alignment likelihoods to estimate the probability that each read originates from a given haplotype, thereby determining the most likely haplotype of origin for each read (Extended Data Fig. 1d). The mutation type (insertion, deletion, or mismatch) was further determined by pairwise alignment of the inferred mutant haplotype to the germline haplotype using pairwise aligner^5^ (v1.81). This refinement alignment incorporated 4-bp flanking sequences on each side.

### Estimating global whole-genome amplification bias

To estimate whole-genome allelic balance for each single cell, we modeled alternative allele counts using a beta-binomial distribution. Germline heterozygous SNPs (hSNPs) were defined as sites with matched-bulk VAF from 0.4 to 0.6 and present in dbSNP. The two shape parameters of the Beta distribution, *α* and *β* were estimated as follows:

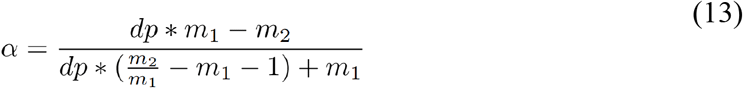

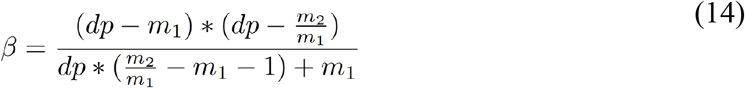

where:

○ *m*_1_ = *mean*(*n*_*alt*_).
○ 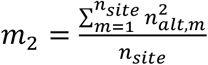
○ *dp* is defined as the median depth of hSNP loci.

Notably, the hSNPs collected for calculating allele imbalance parameters were all of the same median depth, such as 20x. *n*_*alt*_ *r*efers to the alternative read count, and *n*_*site*_ refers to the total number of hSNP loci.

### Estimating local amplification bias using a piecewise gaussian process regression

In whole-genome amplification, uneven amplification across DNA fragments creates locally correlated allelic bias. We addressed this by segmenting reads into fragments using an HMM and then applying piecewise Gaussian process regression to quantify bias within each segment.

To model the correlation of allelic balance, we implemented a piecewise Gaussian process regression framework. First, reads within individual cells were partitioned into independent segments using a hidden Markov model (HMM). Within each segment, the haploid allelic fraction (AF_h_) values of the same haplotype h, derived from phased hSNPs (phased with Eagle2 as described previously), along with their genomic coordinates, served as inputs for the regression. The Gaussian process is formally defined as:

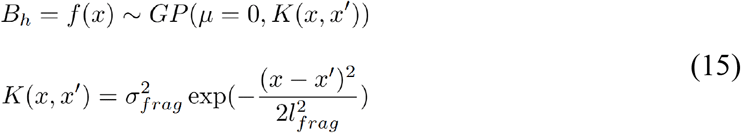

and the allele fraction is represented using a logistic transform:

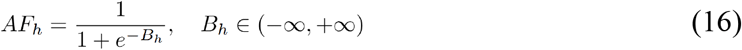

Where:

○ allele imbalance *B*_*h*_ given hSNPs is modelled by Gaussian process *GP* with zero mean function.
○ the radial basis function *K*(*x, x*^’^) with parameter *σ* and *l* ac*c*ounting for fragments attribution.

To prevent discontinuities in the regression line caused by allele dropout, the haploid allelic fraction (AF_h_) was constrained to the interval (10^−7^, 1 – 10^−7^). This model was subsequently applied to predict AF values at each candidate STR loci. Genomic regions containing phased SNPs were excluded if their correlation curves showed sharp fluctuations, specifically those with a mean squared error (MSE) exceeding 0.5, to filter out intervals with potential phasing errors.

### Calculation of mosaic genotyping likelihood with bulk sequencing data

We made no specific assumption about the allele frequencies of mosaic mutations; instead, we adopted a uniform prior distribution over the interval [0, 1]. Based on this, the mosaic genotyping likelihood given bulk sequencing data can be calculated with the following formula:

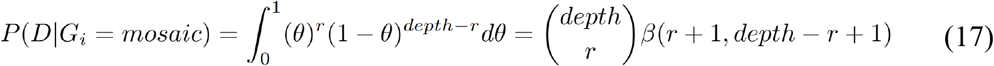

Where:

○ *r* indicates alternative read *c*ount in the bulk sequencing data.
○ *depth* indicates total read depth in the bulk seq data.
○ *θ* indicates the prior distribution of variant allele frequency of mosaic mutations.

### Joint Genotyping

Based on population allele frequencies, sequencing reads and stutter error profiles, BayesMonSTR calculated genotyping posteriors of candidate mosaic STR mutations with a haplotype-based EM-algorithm.

#### Parameter initialization

Population frequencies *f*a were set to be with equal probability. Haplotype phasing probability *P*(*j* − *h*_1_, *k* − *h*_2_|*G*_*i*_) was set to 0.5. *μ*_*msi*_ was set to 0.001. Stutter error parameters *u, d* and *ρ* for each site were calculated with previous procedures.

#### E-step

BayesMonSTR employed a hierarchical Bayesian network to incorporate population, bulk and single cell-level data. We utilized an EM algorithm to calculate maximum likelihood of latent parameters.

Similar to the length-based EM procedure, the E-step for estimating individual germline genotypes at sequence-level resolution was performed as follows:

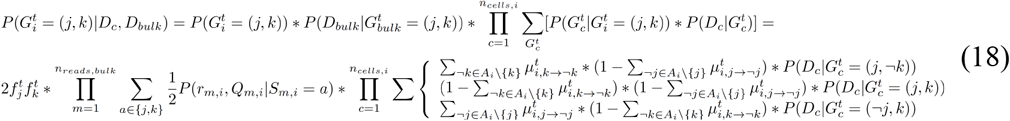

If the germline genotype is homozygous, the prior probability of *G*_*i*_ is 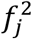 instead of 2*f*_*j*_*f*_*k*_.

Equation 2 and 19 were used for unphased and phased genotype likelihood calculation, respectively.

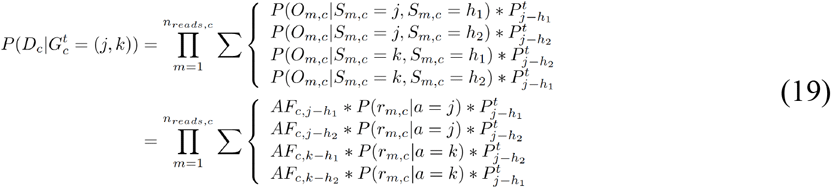

Where:

#### M-step

○ 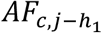 denotes the predicted allele fraction based on STR allele *j* at the locus for cell *c*.
○ 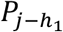 denotes the phasing probability that STR allele *j* is linked with germline hSNP *h*_1_.
○ *P*(*r*_*m,c*_|*a* = *j*) denotes the read likelihoods given allele *j* based on the stutter error model.

#### M-step

We calculated the phasing probabilities that quantify the phasing relationships between STR loci and hSNPs, which was initialized with 0.5. If STR allele *j* linked with germline SNP *h*_2_ and allele *k* links with germline SNP *h*_1_, the phasing probability given individual *i* read information *D*_*i*_ was calculated with the following equation:

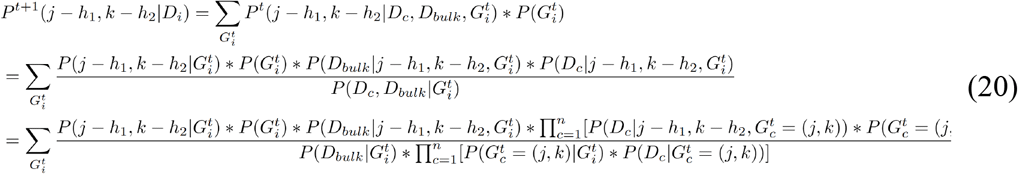

Where:

○ 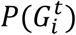 denotes the population haplotype frequency prior.
○ *p*(*D* | *j* − *h*_1_, *k* − *h*_2_) is the genotype likelihood for individual *i* given allele *j* linked with SNP *h*_1_ and allele *k* links with SNP *h*_2_, and with 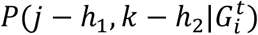 set to 0.5.
○ 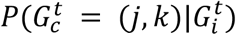 denotes the mosaic mutation probability that the cell genotype is (*j, k*) given *G*_*i*_.

Similarly, the probability that STR allele *j* is linked with germline SNP *h*_2_ follows the same approach. Of note, we only used reads covering both the candidate locus and the nearby hSNP to calculate the phasing probability.

The mosaic allele fraction for mutant allele ¬k was calculated using the following function:

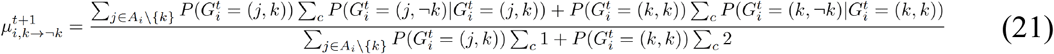

Where:

○ 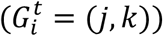 denotes the individual germline probability in E-step.
○ *k* denotes the germline allele.

The sequence-based STR allele population frequency was calculated as follows:

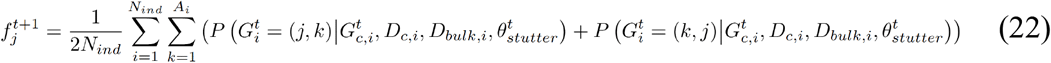

Where:

○ All parameters are defined in Equation (18).

### Estimating somatic genotyping posteriors for each site across cells

BayesMonSTR employed a dynamic programming approach to compute the posterior probability that a candidate locus contains at least one mutant cell. The posterior probability that a mosaic mutation occurs (*k* > ¬k) is calculated with the following formula:

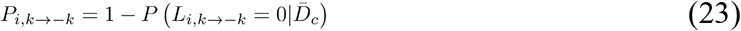

Where:

○ *L*_*i*_ denotes the mutant allele count for individual *i*.

The calculation follows with Bayes’ rule:

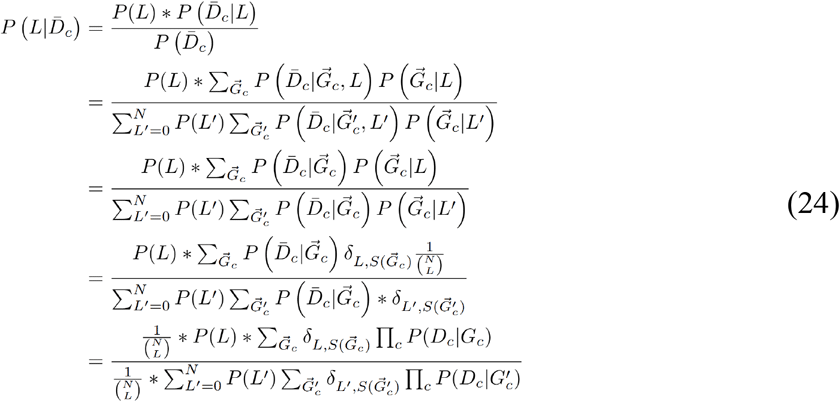

The Kronecker delta function, denoted as *δ*_*L,s*_ is a mathematical function that evaluates to 1 when its indices are equal, and evaluates to 0 when its indices differ.

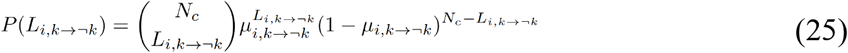

The recursive formula of dynamic programming is as follows:

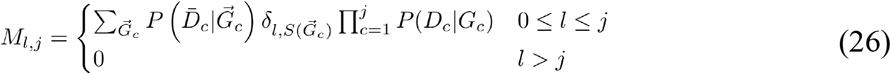

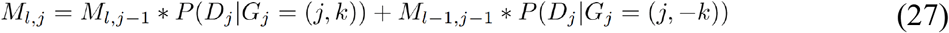

Where:

○ *M*_*l,j*_ denotes probability for mutant allele count *l* and allele *j* in the DP matrix.

### Haplotype phasing related artifacts

Haplotype phasing is widely adopted as a methodology to increase confidence of mosaic mutation calling^6^. The probabilistic haplotype phasing approach we designed is based on two key assumptions: 1) Given a relatively low mutation rate, it is highly unlikely that somatic mutations would occur independently on both haplotypes at the same position; 2) non-cancer cells possess diploid genomes.

Assuming STR allele *j* is linked with hSNP *h*_1_, and STR allele *k* is linked with hSNP *h*_2_. Discordant rate of reads for bulk sample *d*_*bulk*_ and single cell sequencing data *d*_*sc*_ were defined as

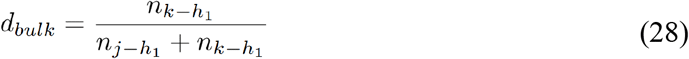

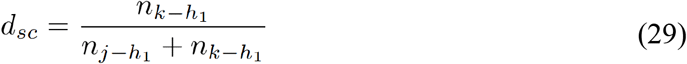

Amplification error *d*_*amp*_ rate and discordant rate *d*_*k*_ for mutant cells sample were defined as:

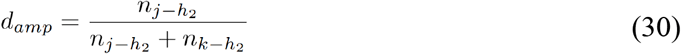

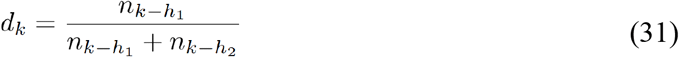

Where:

○ 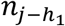 is the number of reads that STR allele *j* is linked with hSNP *h*_1_.
○ 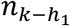 is the number of reads that STR allele *k* is linked with hSNP *h*_1_.
○ 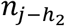 is the number of reads that STR allele *j* is linked with hSNP *h*_2_.
○ 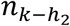 is the number of reads that STR allele *k* is linked with hSNP *h*_2_.

Based on empirical assessment, candidate mutations are considered high-confidence somatic events when all four of their phasing error rates (defined in Equations 28–31) are below 0.1.

### Genome-wide variant prediction using Random Forest (RF) classification model

We trained two Random Forest classifiers to predict mosaic variants using a rigorously curated training set with phasing information (Supplementary Table 1). The first model incorporated phasing-related features (Supplementary Table 1), while the second model was designed for genome-wide prediction, regardless of phasability (Supplementry Table 1). A leave-one-donor-out strategy was applied in model training and testing.

The model was specifically trained to predict refined genotypes, categorizing sites into five distinct classifications: amplification error (amp error), sequencing error (seq error), repeat, heterozygous (het) and mosaic. To ensure an unbiased performance assessment, we employed a leave-one-out cross-validation strategy for model training and testing. To identify mosaic loci, we applied a dual-mode filtering strategy based on phasing feasibility. Phase-informed mode: Variants initially predicted by the phase Random Forest model underwent a series of phasing-based hard filters to yield a high-confidence set of phasing-corrected mosaic loci. These filters included thresholds for bulk, single-cell, and mutant-cell discordance rates, alongside an amplification error rate (all < 0.1, as defined in Equations 28–31), supplemented by requirements for mutant cell VAF ≥ 0.1 and a mutant read count ≥ 2. Non-phaseable mode: For variants lacking reliable phase information, two sequential Random Forest models were applied to mitigate amplification errors. To derive a stringent set of true mosaic sites, we then applied an additional empirical filter requiring mutant cell VAF > 0.25 and mutant read count ≥ 3.

### Examination of artifact sites identified by different software tools

To investigate the superior performance of BayesMonSTR over HipSTR and ExpansionHunter in detecting mosaic STR mutations from single-cell data, we systematically analyzed the false-positive calls from the latter two tools. This analysis revealed how BayesMonSTR’s multi-step filtering pipeline eliminates such artifacts. The key filtering steps are defined as follows: 1. Prescan: Candidate loci lacking alternative alleles or residing in low-depth regions (mean depth ≤ 1 across cells) were removed. 2. Genotyping: Candidate mutations with a mosaic genotyping posterior probability below 0.9 were excluded. 3. Phasing: Candidate mutations were filtered based on the phasing metrics defined in Equations 28–31. Specifically, we excluded any mutation with a bulk discordant rate, single-cell discordant rate, amplification error rate, or mutant-cell discordant rate exceeding 0.1. 4. RF model: Candidate mutations were classified as mosaic if the highest predicted probability was for the mosaic class. The SHAP^7^ (SHapley Additive exPlanations, SHAP, v0.46.0) method was then used to identify the key features driving the RF classification for these mutations (Extended Data Fig. 4e).

### Simulation of spike-in mosaic STR mutations in single cell sequencing data

To simulate mosaic STR mutations in single cell sequencing data, STR indels and mismatches were randomly introduced at target loci using BAMSurgeon^8^ (v1.4.1). Diploid single-cell data were then sampled via beta-binomial sampling to emulate allele-specific amplification imbalance. Finally, aligned reads from 10 single-cell libraries (male donors) were down-sampled and merged to generate synthetic diploid datasets for validation.

### Breast cancer sample collection

A fresh breast cancer tissue sample and a matched normal breast tissue were obtained from a female breast cancer patient at The Second Affiliated Hospital Zhejiang University School of Medicine. The study was approved by the Institutional Review Board of Westlake University (IRB: 20230213DYM001). Informed consent was obtained from the patient in accordance with the IRB-approved protocol, and consent for data publication was also provided. At the time of collection, pathologist analysed the samples and categorized them into tumor and matched normal breast tissue blocks. Fresh tissue samples were transferred to sterile PBS buffer (Sangon Biotech, E607008) on ice and processed immediately upon arrival at the laboratory. The breast cancer and normal tissue samples were then dissociated, and viable single cells were isolated. Single-cell whole genome amplification, library preparation, and whole-genome sequencing were subsequently performed as described below.

### Isolation of single cells of breast tissue samples for scWGS

Fresh tumor and matched normal breast tissue samples were collected and processed within two hours post-surgery. For each sample, small tissue blocks of approximately 1 cm^3^ were isolated for dissociation. The tissue dissociation procedure was performed according to previously described methods^9,10^, with slight modifications. Briefly, tissue samples were manually minced in 1.5 mL tubes containing collagenase type 3 (300 U/mL) (Worthington Biochemical, #LS004182) and hyaluronidase (100 U/mL) (Worthington Biochemical, #LS002592) for 2 hours, with gentle pipetting every 60 minutes. The digested tissue was then centrifuged at 1500 rpm for 5 minutes. The pellet was digested with 5 mL of pre-warmed 0.25% trypsin-EDTA (Thermo #25200072) at 37°C for 5 minutes. Digested cells were washed once with HBSS + 2% FBS (Jackson, #017-000-121), then treated with 0.1 mg/mL DNase I (Worthington Biochemical, #LS002139) in DMEM/F12 media at 37°C for 5 minutes. After washing, the cells were filtered through a 40 μm mesh. Dissociated cells were counted and resuspended in the designated medium for subsequent staining and FACS sorting. Meanwhile, tissue surrounding the small blocks (within 0.5 cm from the edge) was similarly dissociated into a single-cell suspension. DNA was then extracted using a DNeasy Blood & Tissue Kit (Qiagen, 69504) from these suspensions for bulk DNA whole genome sequencing.

For FACS sorting, single cells were stained with the following antibodies: CD45 (BioLegend, 304022), CD31 (BioLegend, 303114), CD49f (Thermo, 12-0495-82), EpCAM (BioLegend, 324204), and CD34 (STEMCELL Technologies, 60013Az.1). The antibodies were used at a dilution of 1:200, with appropriate conjugated fluorophores, and incubated for 30 minutes on ice. After staining, cells were washed and resuspended in Hanks Balanced Salt Solution (HBSS) + 2% FBS + Antibiotic-Antimycotic + 4′,6-diamidino-2-phenylindole (DAPI) (1 µg/mL) at a density of 5 million cells/mL. The stained samples were then analyzed and sorted on a FACSAria Fusion (BD Biosciences) with a 100-µm nozzle. The target single cells were sorted into individual tubes, each containing a single cell and 3 µL of cell buffer, prepared for subsequent single-cell whole genome amplification. The FACS data were further processed using FlowJo V10. Gating strategy is shown in Extended Data Fig. 4.

### Whole genome amplification and single cell DNA & RNA sequencing

The sorted single cells were subjected to whole-genome amplification using either REPLI-G (Qiagen, 150023) or ResolveOME (Bioskyrb). Single cells amplified with ResolveOME yielded both whole-genome amplification products and full-length transcriptome amplification products. In some cases, both reactions were combined to generate both whole-genome amplification products (via REPLI-G) and full-length transcriptome amplification products from the same single cell. All whole-genome and full-length transcriptome amplification products from ResolveOME underwent library preparation using the ResolveOME library preparation protocol, including a 9-10 cycle PCR library amplification. For REPLI-G whole-genome amplification products, as well as the extracted paired bulk DNA, PCR-free library preparation was performed using the Hieff NGS Ultima Pro PCR Free DNA Library Prep Kit V2 (Yeashen, 12196).

The WGS libraries were sequenced on the BGISEQ DNBSEQ-T7 platform (BGI Inc.) using a paired-end sequencing length of 150 bp (PE150). The average sequencing depth for each WGS library was 30–40x, while the bulk WGS library had an average sequencing depth of 150x. Full-length transcriptome libraries were sequenced on the NovaSeq X Plus platform (Illumina Inc.) using the PE150 strategy, generating approximately 20 million reads per library. Additionally, parallel library preparation of the same whole-genome amplification product or parallel sequencing of the same library was conducted to obtain WGS data duplication.

### Processing of RNA sequencing data

For RNA sequencing data, trimmomatic^11^ (v 0.39) was first used to remove adapters. Salmon^12^ (v 1.10.2) was used to quantify the expression of transcripts. The salmon index was constructed using the GTF and FASTA files (v GRCh37.87) from ENSEMBL.

### Comparison of algorithms detecting mosaic STR mutations from the benchmarking dataset

To adapt the two germline mutation callers for identifying mosaic STRs from single cell sequencing data, we applied a series of stringent filters to each software to generate a mosaic mutation call set.

HipSTR^3^ (v0.6.2) was run using the mode of “*de novo* stutter estimation + STR calling with *de novo* allele generation”, variant calls were then filtered following recommended empirical cutoffs (--min-call-qual 0.9 --max-call-flank-indel 0.15 --max-call-stutter 0.15 --min-call-allele-bias -2 --min-call-strand-bias -2). ExpansionHunter^13^ (v5.0.0) with default settings were used for variant callings of benchmark datasets.

Germline genotypes and mosaic genotypes were identified as the most frequent and second most frequent genotypes across all cells. Cell-specific variants were identified by those that have one mutant cell, and shared variants were identified as those with multiple mutant cells. In addition, if there are less than five valid cell calls, the candidate loci were excluded from the mutation list.

For BayesMonSTR, variants were identified using a genotyping cutoff of mosaic posterior ≥ 0.9, predicted as mosaic by the RF model, and passing a series of hard filters. More specifically, for phasable sites, the hard filters include a series of phasing-related filters (equations 28-31, bulk discordant rate < 0.1, single cell discordant rate < 0.1, amplification error rate < 0.1 and mutant cell discordant rate < 0.1), and other filters (VAF ≥ 0.1 and mutant read count ≥2); For non-phasable sites, the hard filters include VAF > 0.25 and mutant read count ≥ 3.

### Cross validation strategies of candidate mutations

To evaluate candidate STR loci detected by different callers, we assessed validation independently at three levels—DNA (duplicate library), RNA (single-cell RNA-sequencing), and bulk sample—using orthogonal evidence derived from the same single cell or its matched bulk sample. DNA-level validation was based on duplicate-library sequencing, RNA-level validation was based on single-cell RNA-sequencing reads, and bulk-level validation leveraged matched bulk sequencing from the same sample. Results from BayesMonSTR_phase, BayesMonSTR_unphase, HipSTR, and ExpansionHunter were first harmonized into a unified framework and then evaluated within each data_type × protocol_type stratum.

For DNA-level validation, a one-sided lower-tail binomial test was applied using the duplicate-library mutant read count (k) and total read count (n). For BayesMonSTR_phase and BayesMonSTR_unphase, two tests were performed, using either the model-derived expected mutant fraction (vaf_mut_sc_mean_prob) or θ = 0.5 as the null probability, and the larger P value was retained for classification. For HipSTR and ExpansionHunter, the binomial test was performed using θ = 0.5 only. For RNA-level validation, a one-sided lower-tail binomial test was applied using the RNA mutant read count (k) and total RNA read count (n), with θ = 0.5 as the null probability. Loci with missing data or zero read depth were classified as undetermined. For the remaining loci, validation outcomes were defined as: validated when P ≥ 0.05 and k > 0; unvalidated when P < 0.05 or k = 0; and undetermined otherwise.

For bulk-level validation, matched bulk sequencing data from the same biological sample were used to assess whether a candidate locus represented a somatic mutation or a germline variant. Two types of bulk data were utilized for this purpose: blood-derived bulk sequencing, serving as a constitutional germline reference, and normal tissue bulk sequencing from the same individual. A locus was classified as germline— and therefore assigned to the unvalidated category, as germline variants and sequencing artifacts do not constitute somatic mutations—if either of the following criteria was met in both the blood and normal bulk samples: (i) the variant allele fraction (VAF) in the bulk sample fell within the range of 0.4–0.6, consistent with heterozygous germline presence, or (ii) a two-tailed binomial test against the null hypothesis of θ = 0.5 yielded P > 0.05, indicating that the allele frequency in bulk was statistically indistinguishable from a germline heterozygous state. Loci not meeting either germline criterion in both bulk samples were retained as candidate somatic mutations for downstream analysis.

When multiple callers reported the same locus within the same data type and amplification protocol but assigned discordant validation states, a predefined caller-priority rule was applied (BayesMonSTR_phase > BayesMonSTR_unphase > HipSTR > ExpansionHunter) to assign a final validation status. DNA-level, RNA-level, and bulk-level results were retained separately; no additional cross-layer priority rule was imposed among duplicate-library, RNA, and bulk evidence. Validation statistics were summarized using unique STR loci as the counting unit. Within each data_type × protocol_type × source stratum, we counted validated, unvalidated, and undetermined loci. The primary validation rate was defined as the number of validated loci divided by the total number of unique loci in that stratum, and, for display purposes, we additionally calculated the fraction of validated loci among clearly classified loci (validated + unvalidated).

Shared variants were further evaluated by manual inspection and PhyloSOLID (v 0.0.1) STR genome validation mode (https://github.com/douymLab/PhyloSOLID). Cell genotype likelihoods along with alternative and total read depth information were used for shared variants evaluation based on phylogenetic tree.

A gold-standard benchmarking dataset was constructed from 29 single cells derived from a breast tissue sample, comprising 7 epithelial cells from tumor tissue (Epi-T), 12 epithelial cells from normal tissue (Epi-N), and 10 immune cells (Imm). Of these, 12 cells had duplicate libraries (sequenced twice), enabling DNA-level validation, and 16 cells underwent both DNA and RNA sequencing from the same cell, enabling RNA-level validation. Single-cell DNA amplification was performed using either PTA or MDA. This dataset served exclusively as a benchmarking resource and is distinct from the public datasets used in cross-validation experiments. To prevent leakage between training and evaluation sets, cell-specific mutations derived from two cells within the benchmarking dataset were used as part of the model training set; these two cells were subsequently excluded from all test and validation analyses, ensuring an unbiased assessment of model performance on the remaining benchmarking samples.

To validate model performance on the public dataset (Supplementary Fig. 5g), we employed a leave-one-out cross-validation strategy. Specifically, when predicting and validating on a given dataset, the model was trained on all remaining datasets excluding the target dataset. For example, predictions for the MDA neuron dataset were generated using a model trained on all other datasets, and similarly, predictions for the single-cell colony skin dataset were generated using a model trained on all datasets except the single-cell colony skin data. This strategy ensures that the model has no prior exposure to the validation target during training, providing an unbiased assessment of generalization performance across independent datasets.

### Visualization of STR mutations

INSIGHT (v 0.0.1) developed by our lab were used for visualization of STR mutations (https://github.com/douymLab/INSIGHT).

### Identification of mosaic SNVs from single cells

Similar strategies were applied to detect mosaic SNVs in the same datasets. After sequence alignment and data preprocessing, raw candidate loci were identified by mpileup in SAMtools^14^ (v1.13) with the parameters ‘-B -d 8000 -q 10 -Q 10’. Then, the mutations were identified by using MosaiSC-DNA^15^ (v0.0.1, https://github.com/douymLab/mosaicSC). The predicted SNVs were further filtered with mutant cell VAF ≥ 0.3 and mutant read count ≥ 3 to get a more reliable set of mosaic SNVs.

### Phylogeny reconstruction using shared variants

Given the modest validation rate for shared variants, the constructed phylogenetic tree (Figure 2g) was based on manually checked cell-by-mutation matrix (Supplementry Table 5). PhyloSOLIDD^16^ (v0.0.1) was used for phylogeny reconstruction.

### Correlation of age with multiple mutation-derived features

We performed association analyses between age and multiple mutation-derived features, including mutation rate, variant length, and cell proportion with >5bp indels. Among these, the mutation rate was calculated by dividing the number of mutations by the total length of callable regions. We used scipy^17^ (v 1.14.1) to perform linear regression analysis between age and features.

### Mutation signature analysis

For mutations in STR regions, mutation matrices were generated using SigProfilerMatrixGenerator^18^ (v 1.2.31), and then, SigProfilerAssignment^19^ (v 0.1.9) was used to assign known COSMIC mutation signatures to every individual, which was accomplished using the forward stagewise algorithm and nonnegative least squares. In our samples, three 83-cat signatures of small insertions and deletions (ID1, ID2, ID12) and multiple 96-cat signatures of single base substitutions were found to be active. Additionally, SigProfilerAssignment was used to assign signature probabilities to every mutation, allowing to quantify the mutation burden associated with each signature for every individual.

Additionally, we also used a redefined InDel taxonomy^20^ to illustrate mutation profile for InDels in STR regions, and the mutation signature similarity was computed via cosine similarity was used to compute similarity between our InDels and those caused by the *MSH2* and *MSH3* knockouts^20^.

We also compared the pattern of STR-region SNVs and the established patterns of extra-STR mosaic SNVs in normal human prefrontal cortex^21^ . The mutation signature similarity was computed via cosine similarity.

Specially, for SNVs in STR regions, a custom method was developed to classify SNVs into different categories. Initially, mutations were distinguished based on the six kinds of fundamental base changes, followed by classification according to the nucleotide context of the entire STR region in which the SNV occurred (considering both repeat unit length and total length of the repeat region). Subsequently, non-negative matrix factorization was performed using MutationalPatterns^22^ (v 3.12.0) to extract three distinct *de novo* mutation signatures. Based on the relative contribution of each signature, the mutation burden associated with each signature for every individual was quantified.

### Relationship between mosaic SNVs in DNA repair gene and mosaic STR indel burden

The list of DNA repair genes was obtained from the comprehensive compilation maintained by the laboratory of Richard D. Wood at The University of Texas MD Anderson Cancer Center (https://www.mdanderson.org/research/departments-labs-institutes/labs/wood-laboratory/resources.html). This curated gene set represents an updated synthesis based on previously published reviews.

ANNOVAR^23^ (v 2020-06-07) was used to identify exonic variants, their associated genes, and their effects on the protein sequence, including synonymous, nonsynonymous, stop-gain, stop-loss, and frameshift variants. Variants annotated as “unknown” by ANNOVAR due to incomplete ORF information were re-annotated using the Ensembl Variant Effect Predictor^24^ (VEP, https://www.ensembl.org/vep) to determine their coding consequences based on updated transcript models.

To assess the association, we fitted a generalized linear mixed model (GLMM) with a negative binomial distribution to account for overdispersion in count data. The model included somatic SNV status (presence in DNA repair gene vs. absence) as a fixed effect and a random intercept for individual to account for within-individual correlations (e.g., multiple neurons or samples per individual). The significance of fixed effects was evaluated using Wald tests. All analyses were performed in R (v 4.3.3) using the glmmTMB^25^ (v 1.1.14) package.

### Identification of potentially functional-disrupting mosaic STR mutations

ANNOVAR was employed to determine the nearest gene of mosaic STR mutations and identify potentially damaging exonic variants, including nonsynonymous, stopgain, stoploss, and frameshift variants. CADD^26^ (v 1.7) scores were incorporated to assess variant deleteriousness. ENCODE’s candidate cis-regulatory elements annotation^27^ was used to identify variants with potential regulatory functions. The chromatin state annotation from ENCODE was used to identify variants in active TSS and enhancer regions. Chromatin state annotations were derived from the ChromHMM 18-state model from the ENCODE portal^28^ for seven primary B cell samples, 17 lung tissues and 16 brain tissues.

Potentially functional-disrupting STR indels were defined as variants in upstream, UTR or exonic regions annotated by ANNOVAR, or regions defined as chromatin state of active TSS and enhancer by ENCODE, or variants have CADD score greater than 20.

Gene affected by a STR indel was determined as the nearest gene annotated by ANNOVAR.

### Identification of mosaic STR mutations in functional non-coding elements

To identify STR mutations may have a significant impact on gene expression, we utilized the human G-quadruplexes prediction panel from EndoQuad^29^ for G-quadruplex forming regions, the Level 2 R-loop annotation panel from R-loop Base^30^ for R-loop forming regions, ENCODE’s candidate cis-regulatory elements annotation^27^ for CTCF-binding sites, and the CpG islands annotation from the UCSC Genome Browser^31^ for CpG islands.

### Identification of mosaic STR mutations near genes of high expression

Gene expression data of whole blood, lung and brain cortex tissues was downloaded from GTEx^32^. Within each tissue, we calculated quartiles of expression of all genes near STR loci and categorized each gene into four categories (Q1 to Q4) based on average expression level across samples, with Q1 representing expression level is in minimum to first quartile.

### Gene ontology (GO) and disease enrichment

KOBAS^33^ (v 3.0) was used to conduct Gene Ontology (GO) and Kyoto Encyclopedia of Genes and Genomes (KEGG) disease enrichment of genes near STR indels. To be specific, Fisher’s exact test was used to calculate P-values, while Benjamini and Hochberg (1995) method was used to control the FDR.

### Process of snATAC-seq data

Raw FASTQ files of snATAC-seq were aligned to the hg38 reference genome, using 10x cellranger-atac count toolkit (v 2.1.0). Only cells identified through cellranger’s cell calling were included in subsequent analyses. Duplicate reads were first filtered to ensure data integrity. Reads sharing identical cell barcodes and genomic coordinates (start and end positions) were grouped. Within each group, only reads with identical sequences were considered duplicates; among these, the read with the highest mapping quality and average base quality was retained.

### Detection of mosaic STR mutations from snATAC-seq

Building upon the framework of BayesMonSTR, we adapted the approach to better handle the sparsity of single-cell ATAC-seq data by performing genotyping at the pseudo-bulk level.

The stutter error model was trained using 26 snATAC-seq datasets derived from brain tissue and whole blood samples, obtained from the publicly available studies SRP309633 and SRP192525. After estimating the stutter error model and extracting alleles through STR segmentation as mentioned before, we directly estimated individual-level mosaic posterior probabilities at the pseudo-bulk level.

We than perform multiple strict filters to retain highly confident mutations including

1. posterior probability exceeding 0.5.
2. count of used reads in genotyping exceeding 10.
3. homozygous in the bulk genotype with exactly one alternative read.
4. no other reads at the locus contained off-target stutter artifacts or mismatches.
5. recurred in less than two individuals.

### Per-cell mutation rate estimation

In snATAC-seq data, the mutation rate per cell can be estimated for each individual cell as the ratio of the number of detected mutations to the size of its callable region. Callable regions were defined as regions covered by at least one read spanning STR with high mapping quality. And the mutation rate of an individual (*μ*) can be estimated by the mean of cells’ mutation rate:

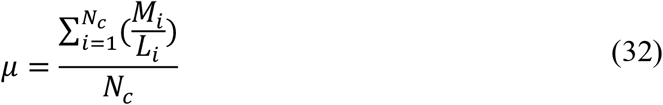

Where:

○ *M*_*i*_: the number of somatic mutations detected in cell *i*;
○ *L*_*i*_: the total length (bp) of the callable genomic regions (i.e., sequenced with sufficient coverage and quality) in cell *i*;
○ *N*_*c*_: the total number of cells analyzed.

The mutation rate of the regulatory region was calculated using the same strategy, with the exception that regulatory callable regions were used instead of whole callable regions.

### Dual-Luciferase promoter reporter assay

The promoter sequences of *NRN1* and *CNTNAP2* were PCR-amplified from HEK293T cDNA templates and cloned into our dual-luciferase reporter plasmid vector. All constructs were verified by sequencing. 0.2M HEK293T cells were seeded into 24-plate well with DMEM for 12 hours, cells were transiently transfected with plasmids using polyethyleneimine transfection reagent (1mg/mL; Polyscience, #23966-1). After 48 hours, promoter activity was analyzed using the Dual-Luciferase Reporter Gene Assay Kit (Yeasen Biotechnology, 11402ES60) following manufacturer’s protocol on a Varioskan LUX Microplate reader (Thermo). Relative promoter activity was calculated as the ratio of firefly/Renilla luminescence.

## Data availability

Bulk WGS data and single cell data from breast cancer samples generated in this study will be publicly available in the Sequence Read Archive (SRA) (PRJNA1378499). WGS data for neurons, B cells, and lung epithelium are accessible through dbGaP under accession numbers phs001485.v3.p1 and phs001808.v1.p1, and in the European Genome-phenome Archive (EGA) under accession EGAD00001005193, respectively. sn-ATAC-seq data from brain tissue are available in the EGA (accession EGAS00001001934) and Synapse (accession syn52293417).

## Code availability

BayesMonSTR is implemented in Python and R and is licensed under the MIT License. The source code, documentation and examples are available on GitHub at https://github.com/douymLab/BayesMonSTR.

## Acknowledgements

We are deeply grateful to the generous tissue contributions. We are grateful to Dr. Peter J. Park for his critical insights and discussions; Drs. Xushen Xiong, Manolis Kellis, Xiaodong Liu, Jing Zhang, Shumin Duan and Dan Zhou for data sharing and discussions; and Drs. Shang Cai, Danyang He, Hongtao Yu, Xiaochun Yu, Wei Wu, Lovelace J. Luquette, Ge Gao, Kai Zhang, Yanxiao Zhang, Jia Zheng, Yuxiang Huang, and Dangsheng Li for their comments and assistance. This work was supported by the Westlake Laboratory of Life Sciences and Biomedicine (Hangzhou 310024, Zhejiang, China), under the grant “Key R&D Program of Zhejiang” (2024SSYS0032) as well as the National Natural Science Foundation of China (32270682) to Y.D.; and the National Natural Science Foundation of China (82525055 and 82473121) to Q.X. We thank the Flow Cytometry Core Facility and the Genomics Core Facility at Westlake University for their experimental assistance, and the High-Performance Computing Center for technical support. We would like to thank the submitters of the following datasets: dbGaP accession phs001808.v1.p1, dbGaP accession phs001485.v3.p1, the European Genome-phenome Archive (EGA) accession EGAD00001005193, the EGA accession EGAS00001001934 and Synapse accession syn52293417. Data used in this study under the dbGaP accession phs001485.v3.p1 was generated by Drs. Christopher A. Walsh and Peter J. Park with funding from NINDS grant R01NS032457. The public datasets were obtained from dbGaP at http://www.ncbi.nlm.nih.gov/gap and the EGA at https://ega-archive.org/.

## Author contribution

Y.D. conceived and supervised the project and secured the funding. Q.X., D.H., X.L., S.C., X.X., M.K. provided support in the supervision of the project. C.W., W.F., W.W., Y.X., J.L., X.M. and J.Y. all made significant contributions to the project. W.F. contributed to software development with help from C.W., W.W., Y.X., J.L. and Y.Z., C.W. developed the pipeline analyzing snATACseq data and lead the bioinformatics analysis. Y.X contributed to phylogeny reconstruction, public data downloading and developed the mutation-visulization software. Q.Y. helped phylogeny reconstruction. J.L. was responsible for generating the benchmarking dataset with help from X.Y., X.M., Y.L., and Z.M., X.Y. performed the breast tumor surgery. J.Y. and X.M. performed the functional assays. N.Y. contributed to cell sorting and related wet lab experiments. Y.Z. and J.L. were responsible for generating non-STR SNV mutation callings. W.L, W.W., C.W., Y.X, W.F. and H.L. contributed to mutation examination. Y.D., C.W., W.F., W.W. and J.L. wrote the manuscript. All authors carefully reviewed and approved the final manuscript. All authors in the Data Group contributed to data access, data generation, or data transfer.

## Inclusion & Ethics statement

This study was conducted in accordance with ethical guidelines and principles. It involved one human participant, and no animal subjects were included. The authors affirm that all research procedures were performed with respect for the participant’s rights and well-being. The sequencing data generated in this study are publicly available, and no personally identifiable information was included. The authors ensure that the study adheres to the highest ethical standards in research, data generation, and usage.

## Declaration of generative AI and AI-assisted technologies

During the preparation of this work the authors used DeepSeek in order to polish our language. After using this tool, the authors reviewed and edited the content as needed and take full responsibility for the content of the published article.

